# Bacterial growth temperature as a horizontally acquired polygenic trait

**DOI:** 10.1101/2024.09.13.612959

**Authors:** Anne A. Farrell, Camilla L. Nesbø, Olga Zhaxybayeva

## Abstract

Evolutionary events leading to organismal preference for a specific growth temperature, as well as genes whose products are needed for a proper function at that temperature, are poorly understood. Using 64 bacteria from phylum *Thermotogota* as a model system, we examined how optimal growth temperature changed throughout *Thermotogota* history. We inferred that *Thermotogota*’s last common ancestor was a thermophile and that some *Thermotogota* evolved the mesophilic and hyperthermophilic lifestyles secondarily. By modeling gain and loss of genes throughout *Thermotogota* history and by reconstructing their phylogenies, we demonstrated that adaptations to lower and higher growth temperature require both the acquisition of necessary genes and loss of unnecessary genes. Via a pangenome-wide association study, we correlated presence/absence of 68 gene families with specific optimal growth temperature intervals. While some of these genes are poorly characterized, most are involved in metabolism of amino acids, nucleotides, carbohydrates, and lipids, as well as in signal transduction and regulation of transcription. Most of the 68 genes have a history of horizontal gene transfer to/from other bacteria and archaea, suggesting that parallel acquisitions of genes likely promote independent adaptations of different *Thermotogota* species to specific growth temperatures.

**Significance:** While the currently known range of life-permitting temperatures spans from −15°C to +122°C, most living organisms, including microbes, can grow only in a narrow temperature interval around their optimal growth temperature. The genetic and genomic determinants of such preference remain poorly understood. Using genomes from *Thermotogota*, a group of bacteria that collectively can grow between 20° and 90°C, we detected 68 genes, presence of which strongly correlates with growth at specific optimal growth temperature. Our findings dramatically expand a list of genes that are likely important for both lowering and increasing preferred growth temperature of a microorganism. We also demonstrated that these genes were usually horizontally acquired from other bacteria and archaea that likely share environment with *Thermotogota*, highlighting the importance of gene exchange in microbial adaptation.

## Introduction

Temperature is one of the most important physical parameters that define the distribution, diversity, and abundance of living organisms (Schumann 2009). The currently known range of life-permitting temperatures spans from −15°C to +122°C (Mykytczuk et al. 2013; Takai et al. 2008), with most of this gamut occupied exclusively by microorganisms. Individual lineages are usually adapted to grow only within a narrow temperature interval, which is likely due to the high evolutionary costs of being a thermal generalist: fluctuations in temperature require many physiological changes, such as alterations to cell wall and membrane composition, translation, and energy metabolism (Mykytczuk et al. 2013; Barria et al. 2013; Phadtare 2004; Pollo et al. 2015). Within the temperature range tolerated by an organism, there is a specific, or optimal, growth temperature (OGT) at which the organism exhibits the fastest growth. Based on their OGT, organisms are broadly classified as psychrophiles (OGT< 15**°**C), mesophiles (15-45**°**C), thermophiles (>45**°**C), and hyperthermophiles (>80**°**C). Disjoint distribution of organismal temperature predilections across the Tree of Life (Pollo et al. 2015) suggests that OGT, as a trait, has evolved multiple times.

The genetic and genomic determinants of the preference to grow at specific temperature remain poorly understood. What genes/proteins/enzymes are needed for life at a particular temperature? To date, only a few proteins have been experimentally shown to be required for growth at specific temperatures. For example, reverse gyrase (Rgy) is required for growth at temperatures exceeding 90°C (Lipscomb et al. 2017) and the tRNA m^1^A58 methyltransferase (TrmI) is required for growth at temperatures above 80°C (Droogmans et al. 2003). How does the evolutionary transition between thermophilic and mesophilic lifestyles happen? For example, in a laboratory evolution of mesophilic *E. coli* into a facultative thermophile, deletion of the glycerol transporter gene (*glpF*) and mutations in a fatty acid desaturase/isomerase gene (*fabA*) contributed to the increase of *E. coli*’s OGT from 37 to 48°C (Blaby et al. 2012). To what extent is such transition influenced by mutation, gene duplication and horizontal gene exchange? The reverse gyrase gene was shown to be acquired horizontally multiple times by unrelated hyperthermophiles (Catchpole & Forterre 2019). Mesophilic bacteria may have evolved from their thermophilic ancestors by horizontal acquisition of genes needed for growth at lower temperatures (Pollo et al. 2015; Zhaxybayeva et al. 2012; López-García et al. 2015), in particular of genes involved in regulation, signal transduction, secondary metabolite biosynthesis, and amino acid metabolism (Zhaxybayeva et al. 2012). Adaptation to growth at low temperature in bacteria has also been associated with the expansion of gene families, especially in psychrophiles (e.g., (Sabath et al. 2013; Piette et al. 2010)).

The bacterial phylum *Thermotogota* represents an excellent model system to further tackle above-raised questions, as it comprises mesophiles, thermophiles, and hyperthermophiles that collectively can grow between 20° and 90°C and have OGTs between 37°C and 80°C (Nesbø et al. 2021; Nesbø, Farrell, et al. 2019). Additionally, *Thermotogota*’*s* small, compact genomes, which range from ∼1.75 Mb in hyperthermophilic *Thermotoga* spp. to ∼2.9-3.2Mb in mesophilic *Mesotoga* spp., reduce the search space for the possible components needed for growth at specific temperature. Via an analysis of 64 *Thermotogota* genomes and 16S rRNA genes of 60 validly described *Thermotogota* species that have experimentally determined OGTs, we inferred changes in OGT from the Last Common *Thermotogota* Ancestor (LCTA) to present-day *Thermotogota*, systematically examined how gene content changes across *Thermotogota* genomes, detected genes whose presence significantly associates with growth at specific OGT, and examined evolutionary histories and functions of these genes.

## Results

### Evolution of OGT in *Thermotogota* follows the evolutionary history of the phylum

First, we evaluated if the evolution of OGT in *Thermotogota* is coupled to the evolutionary history of their genomes, or if there are other selective forces at play. Using a phylogeny of concatenated ribosomal proteins from 64 *Thermotogota* genomes as a reference tree and OGT of the corresponding species as trait values (**Supplemental Table S1**), we found that evolution of OGT is strongly correlated with phylogeny (Pagel’s lambda = 0.93; p << 0.001). The correlation is consistent with the OGT changes across *Thermotogota* occurring under a random walk, Brownian motion model (Felsenstein 1973). Further model testing revealed that an Ornstein-Uhlenbeck model (Felsenstein 1988) - a Brownian motion model in which the variance of the trait is constrained - has a better fit than the classic Brownian motion model (AICw = 0.99, **Supplemental Table S2**). Our finding suggests that as *Thermotogota* diversified, their OGT co-evolved with their genomes. Thus, we can use the evolutionary history of the phylum to infer and track OGT changes from the LCTA to extant *Thermotogota*.

### Modeling OGT predicts a moderately thermophilic common ancestor of *Thermotogota* and multiple shifts in OGT within *Thermotogota*

We modeled evolution of OGT in *Thermotogota*, as it diversified from LCTA, using three independent approaches. First, we modeled OGT as a continuous trait that changes along the reference phylogeny under a Bayesian random walk (**Figure 1**). This model predicted that the LCTA was a thermophile, with its mean OGT at 65.25°C (95% CI of 42.71°C-88.15°C from 80,000 estimates).

**Figure 1.**
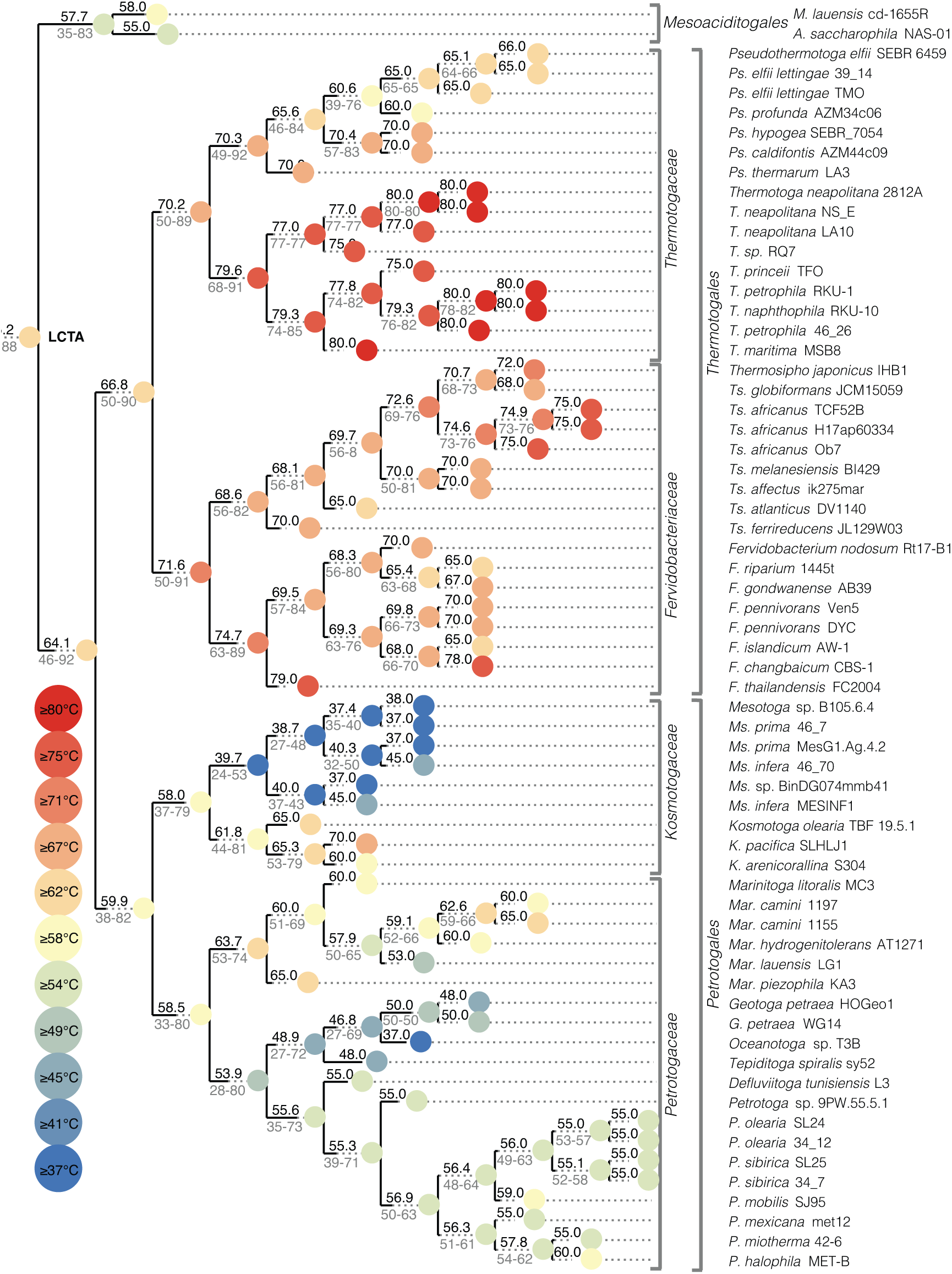
Predicted change in OGT from the LCTA to 64 present-day *Thermotogota* as inferred using Bayesian random walk model. The OGT values are mapped onto the rooted reference phylogeny reconstructed using 50 concatenated ribosomal proteins. Outgroup is not shown. Nodes are colored by either estimated (internal nodes) or extant (external nodes) OGT (in °C). For each internal node, estimated mean OGT is shown in black font above the parent branch, while rounded 95% credible interval is given in grey font below the branch. Scale bar, substitutions per site.

Second, we inferred OGT using the GC-content of 16S rRNA stems, which robustly correlates with OGT across bacteria (Hu et al. 2022; Hildebrand et al. 2010; Musto et al. 2006). In *Thermotogota*, we evaluated the correlation using two datasets: the 64 representative *Thermotogota* strains for which genomes are available and the 60 validly described *Thermotogota* species with experimentally determined OGT. The latter dataset contains 12 species that are not represented among the 64 *Thermotogota* strains due to lack of publicly available sequenced genomes (see **Methods** for details of the dataset construction). For both datasets, we observed a moderately strong correlation between GC content of 16S rRNA stems (*F*GC) and their OGTs (*T*opt = −78.89 + 197.65*F*GC, MSE = 47.1, R^2^=0.65 for the “64 *Thermotogota* strains” dataset and Topt = −72.32 + 189.19 *F*GC, MSE = 30.64, R^2^ = 0.72 for the “60 *Thermotogota* species” dataset; **Supplemental Figure S1**). For the “60 *Thermotogota* species” dataset, the correlation remains statistically significant independent of the phylogenetic signal (phylogenetic contrasts analysis; R^2^ = 0.1, p-value = 0.00953; **Supplemental Figure S1**).

In the analyses of the “60 *Thermotogota* species*”* dataset under a homogeneous model, the stems from the reconstructed 16S rRNA of the LCTA have a GC content of 73.3% and a inferred LCTA’s OGT of 66.36 ± 1.47°C (**Supplemental Figure S2**). The analyses of the “64 *Thermotogota* strains” dataset under the same model gave nearly identical OGT estimate of 66.58 ±1.87°C (**Supplemental Figure S3**). In the analyses of the “60 *Thermotogota* species*”* dataset using a non-homogeneous model, which accounts for potential composition shifts along different branches in the *Thermotogota* tree (Yang 2007), the stems from the reconstructed 16S rRNA of LCTA have a slightly higher GC content of 74.4%, and the inferred LCTA’s OGT of 68.49 ± 1.58°C. The analyses of the “64 *Thermotogota* strains” dataset under the same model yielded an OGT estimate of 64.21 ± 1.43°C.

In our third approach, we examined amino acid composition of encoded proteins, which also exhibits a strong correlation with OGT across prokaryotes (Sauer & Wang 2019; Zeldovich et al. 2007; Sun et al. 2020). Specifically, median fraction of the “IVYWRELGKP” amino acids across all non-transmembrane proteins encoded in a genome (*F*IVYWRELGKP) correlates very strongly with OGT (Sauer & Wang 2019), and this correlation is present in our dataset of 64 *Thermotogota* genomes (Topt = −403.39 + 754.81 *F*IVYWRELGKP, MSE = 47.25, R^2^ = 0.67). For a subset of 294 non-transmembrane proteins broadly represented in 64 *Thermotogota* genomes, the median IVYWRELGKP of the reconstructed ancestral proteins of each gene family’s last common ancestor is 0.621, which corresponds to a 65.82 ± 1.79°C estimate of the LCTA’s OGT.

In all three approaches, the mean OGT of the LCTA is lower than the previously reported estimate of ∼76°C (Green et al. 2013). Within *Thermotogota*, there are multiple shifts in OGT towards both higher and lower OGTs (**Figure 1**; **Supplemental Figures S2** and **S3**), suggesting many independent adaptations of *Thermotogota* lineages to different temperature conditions.

Particularly notable, and consistent across inferences using all three approaches, are the OGT decreases on the branches leading to the last common ancestors of the *Petrotogales* and *Mesoaciditogales* orders, further OGT decreases in *Mesotoga* species and increases in *Thermotoga* species.

### Associations between changes in genome content and changes in OGT

Genome size correlates negatively with OGT in microbes (Sauer & Wang 2019; Sabath et al. 2013), with thermophiles generally having smaller genomes than mesophiles. Across 64 *Thermotogota*, the correlation is modest but significant (R^2^ = 0.31; p = 1.82e-06). The hyperthermophiles in the *Thermotogales* order have the smallest genomes (less than 2 Mbp), while the species with the lowest OGTs, *Mesotoga* spp., have the largest genomes (up to 3 Mbp) (Zhaxybayeva et al. 2012; Pollo et al. 2015; Nesbø, Charchuk, et al. 2019). For *Mesotoga* spp., genome size increase is due to both the larger intergenic regions and bigger sizes of gene families (Zhaxybayeva et al. 2012). To examine how genome content has evolved in *Thermotogota* since their diversification from the LCTA and whether any major genome content changes could be correlated with the estimated shifts in OGT, we modeled gain-loss-expansion-reduction of gene families using the presence/absence of each gene family across *Thermotogota* and compared these events to the OGT estimates from the Bayesian random walk model. While gains and losses of genes occur throughout *Thermotogota*, it is evident that the branches with the largest number of gains and losses encompass taxa with OGTs that are either substantially lower or substantially higher than the estimated OGT of LCTA (**Figure 2**). Specifically, the largest number of gene gains are predicted to have occurred on the branches leading to taxa with lower OGTs: 317 gains and 220 losses in the last common ancestor of *Kosmotogaceae*, 552 gains and 130 losses in the last common ancestor of *Mesotoga*, 255 gains and 82 losses on the branch leading to *Oceanotoga* sp. T3B, 200 gains and 98 losses on the branch leading to *Tepiditoga spiralis*. However, large genome content changes are not restricted to the low-temperature *Thermotogota*. The branch leading to hyperthermophilic *Thermotoga* species is estimated to have had 243 gains and 171 losses. The clade of *Marinitoga* spp., which generally have temperatures higher than their *Petrotogaceae* relatives, is predicted to gain 229 genes. The temperature generalist *Kosmotoga olearia* is inferred to have 177 gains and 147 losses.

**Figure 2.**
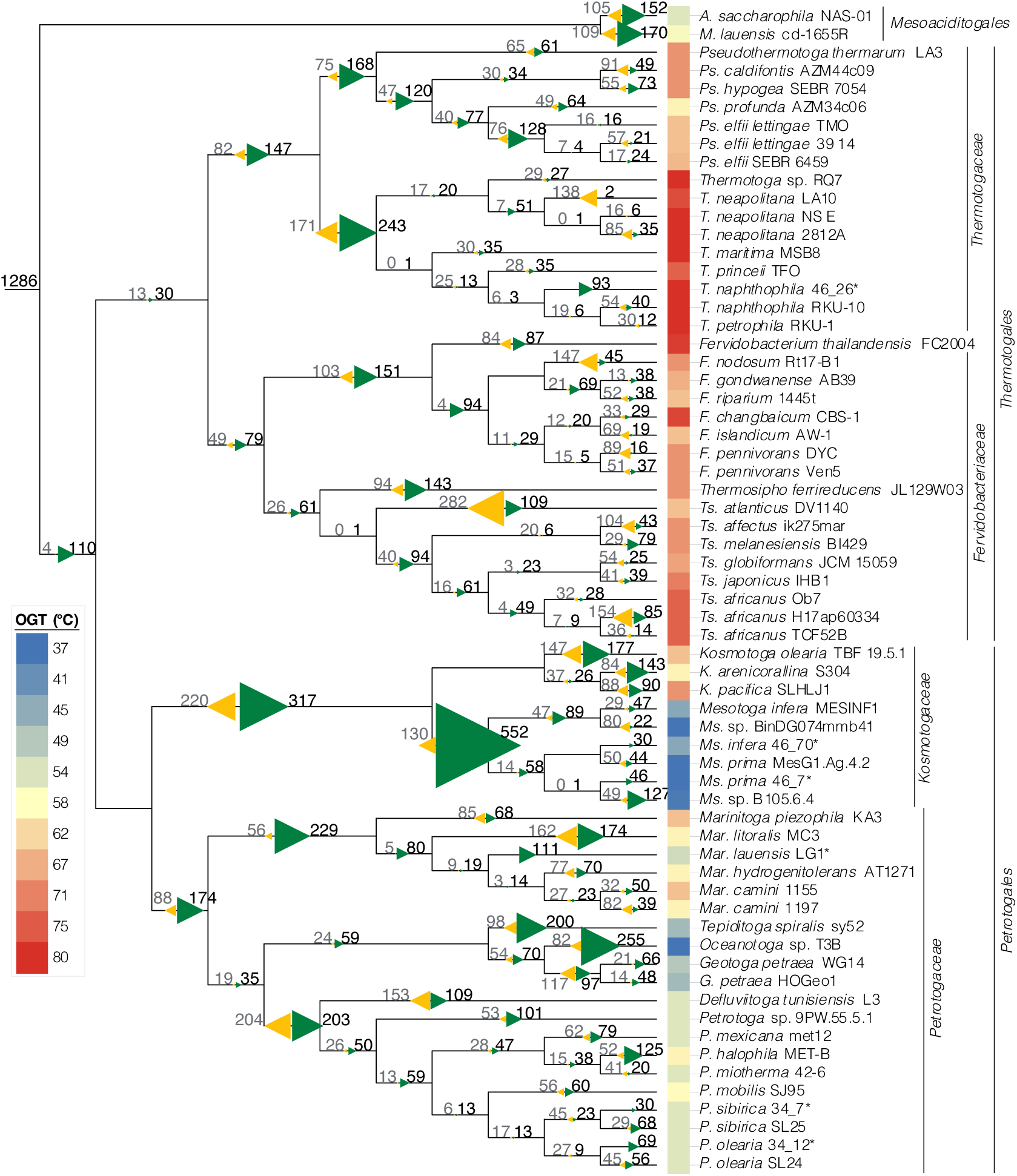
Gains and losses of gene families from the LCTA to the extant *Thermotogota*. Gains and losses modeled with the COUNT program are mapped to the cladogram of the rooted reference phylogeny of 64 *Thermotogota* (the outgroup not shown). At each branch, gains are represented by numbers in black and green right-pointing triangles, while losses are shown by numbers in grey and yellow left-pointing triangles. Only predictions with a posterior probability above 0.6 are included. On branches leading to taxa with incomplete genomes (<90%, marked with an asterisk) the loss values are not shown, since incomplete genomes can cause inflated loss values. OGT of the extant *Thermotogota* is depicted by the color squares next to the strain name. Scale bar, substitutions per site.

### Presence-absence of 68 *Thermotogota* gene families correlates with the different optimum growth temperatures

To evaluate which of the genome content change is associated with shifts in OGT, we conducted a pangenome-wide association study of the 3,604 unique gene presence-absence (phyletic) patterns for correlation with OGT, as implemented in Pyseer (Lees et al. 2018). One hundred twenty-five phyletic patterns, spanning 597 gene families, correlate significantly with OGT (p < 1.42E-05). We narrowed down this list to 68 gene families, whose presence most strongly correlates with optimal growth in one of the three specific temperature intervals: mesophilic growth at 37-45°C (29 gene families), thermophilic growth at 48-59°C (9 gene families), and thermophilic growth at 60-80°C (30 gene families) (**Supplemental Figures S4-S6,** and **Supplemental Tables S3-S5;** see **Methods** for selection criteria).

Based on predictions of a duplication-transfer-loss (DTL) model (Morel et al. 2020), only 19 of the 68 gene families (28%) were present in the LCTA, while the remaining 49 appeared later in the *Thermotogota* history (**Supplemental Figure S7**). Consistent with the earlier discussed modeling of gain-loss-expansion-reduction of gene families, no gene families were gained on the branch leading to *Thermosipho* and *Fervidobacterium*, the two genera that have OGTs similar to the predicted OGT of LCTA. Thirteen of the 19 families present in LCTA and 48 of the 49 families that originated later are collectively predicted to have experienced 293 gene transfer events, highlighting the importance of horizontal gene transfer (HGT) in temperature adaptation of *Thermotogota*. It is well documented that *Thermotogota* genomes undergo extensive HGT with taxa outside the phylum, such as with the *Bacillota* (formerly *Firmicutes*) and Archaea (Nelson et al. 1999; Swithers et al. 2012; Zhaxybayeva et al. 2009), thus opening a possibility that some of the 68 gene families were horizontally transferred to/from *Thermotogota*. By expanding the gene families to include homologous sequences from taxa outside the *Thermotogota* phylum, we found that in 46 of the 63 expanded gene trees with homologs outside of *Thermotogota* (73%), *Thermotogota* do not form a group separately from other taxa (**Figure 3**), further corroborating that the temperature-associated genes were likely acquired and/or donated by *Thermotogota* throughout its evolutionary history.

**Figure 3.**
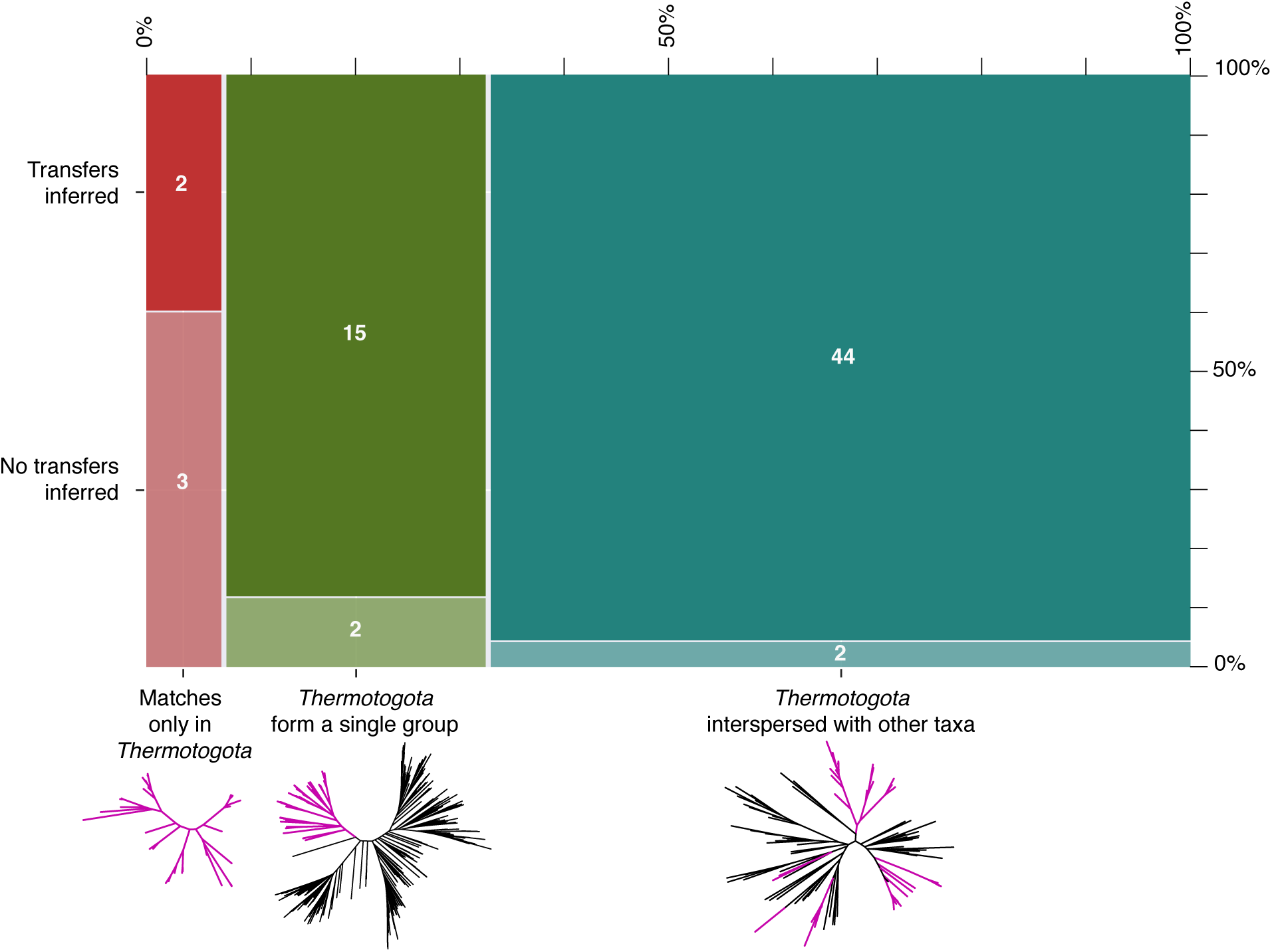
Relationship between the inferences of HGT by the DTL model and the position of *Thermotogota* on the expanded gene family phylogenies. The 68 gene families are categorized based on whether they had no matches in taxa other than *Thermotogota* phylum (in red, 5 families), *Thermotogota* formed a group separate from other taxa (in green, 17 families), or *Thermotogota* interspersed among other taxa (in teal, 46 families). An example tree for each topology category is shown, with branches leading to *Thermotogota* colored in magenta. Each category is further subdivided into those with (solid color) or without (translucent color) transfers, as inferred by the DTL model.

The majority of the 68 families are either localize to the cytoplasm (68%) or are membrane bound (28%) (**Supplemental Figure S8** and **Supplemental Tables S3-S5**), although this is not significantly different from the overall fraction of these localizations among all gene families. Of 60 families that can be assigned to the “Cluster of Orthologous Genes” (COG; (Galperin et al. 2021)) functional categories and of 37 that have Kyoto Encyclopedia of Genes and Genomes (KEGG; (Kanehisa & Goto 2000)) terms associated with a KEGG Pathway, many are involved in transport and metabolism of amino acids, carbohydrates and inorganic ions, as well as in energy production and metabolism (**Figure 4** and **Supplemental Tables S3-S5**). This suggests that different enzymes and different transporters across membrane might be needed for metabolisms at different growth temperatures. Seventeen of the 68 gene families are assigned to “general function prediction only” (COG category R), “function unknown” (COG category S), or not assigned to any COG category or KEGG pathway (**Supplemental Tables S3-S5**), highlighting that a few gene families potentially important for growth at specific OGT are yet to be characterized. Below, we highlight 23 gene families, whose presence correlates best with growth at particular temperature intervals (**Figure 5** and **Table 1**) and evaluate the genes’ possible origins and evolutionary trajectories.

**Figure 4.**
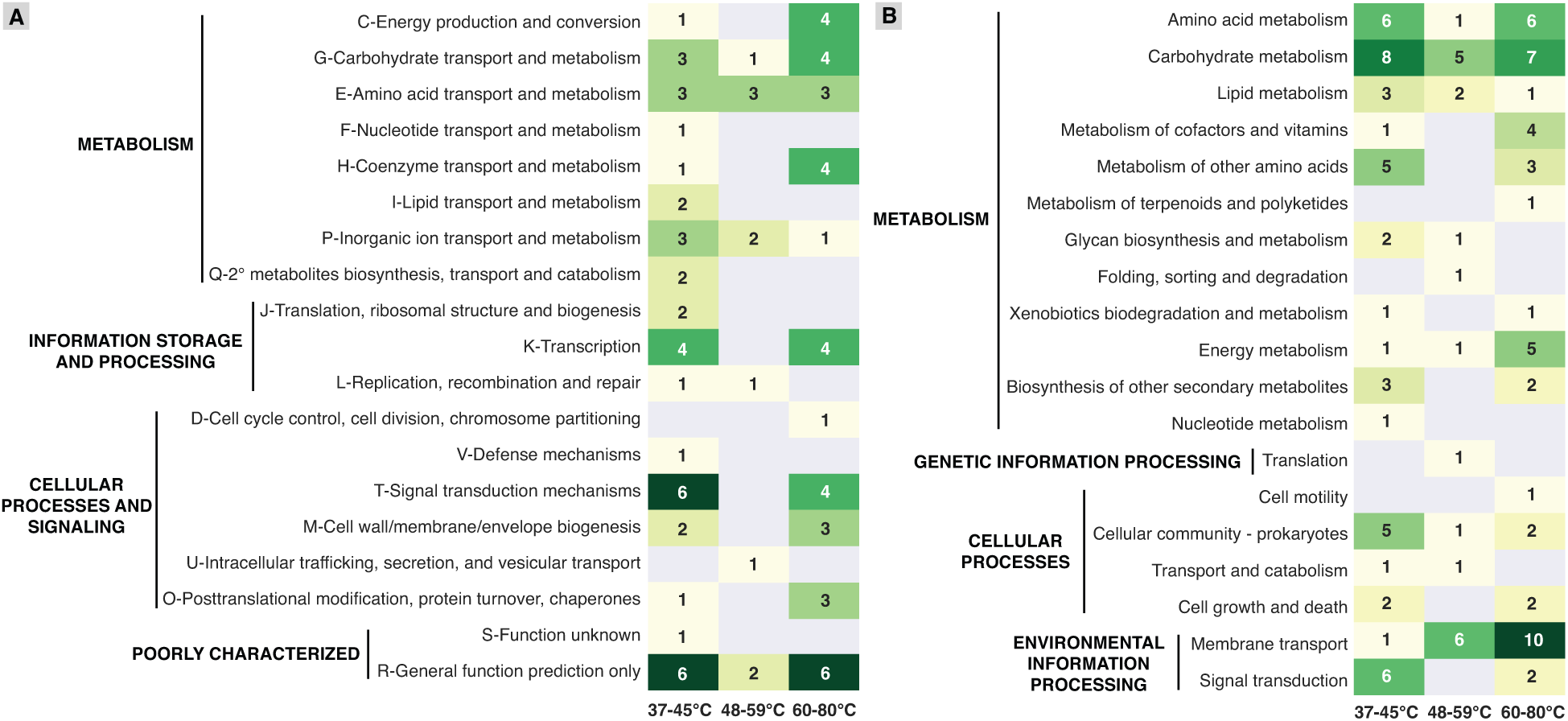
Distribution of 60 gene families with assigned COG categories (panel A) and 37 gene families with assigned KEGG pathways (panel B) among the three temperature intervals. Each cell lists the number of gene families. Families that were assigned to more than one category are listed in each category. Description of each COG category is preceded by its one-letter abbreviation.

**Figure 5.**
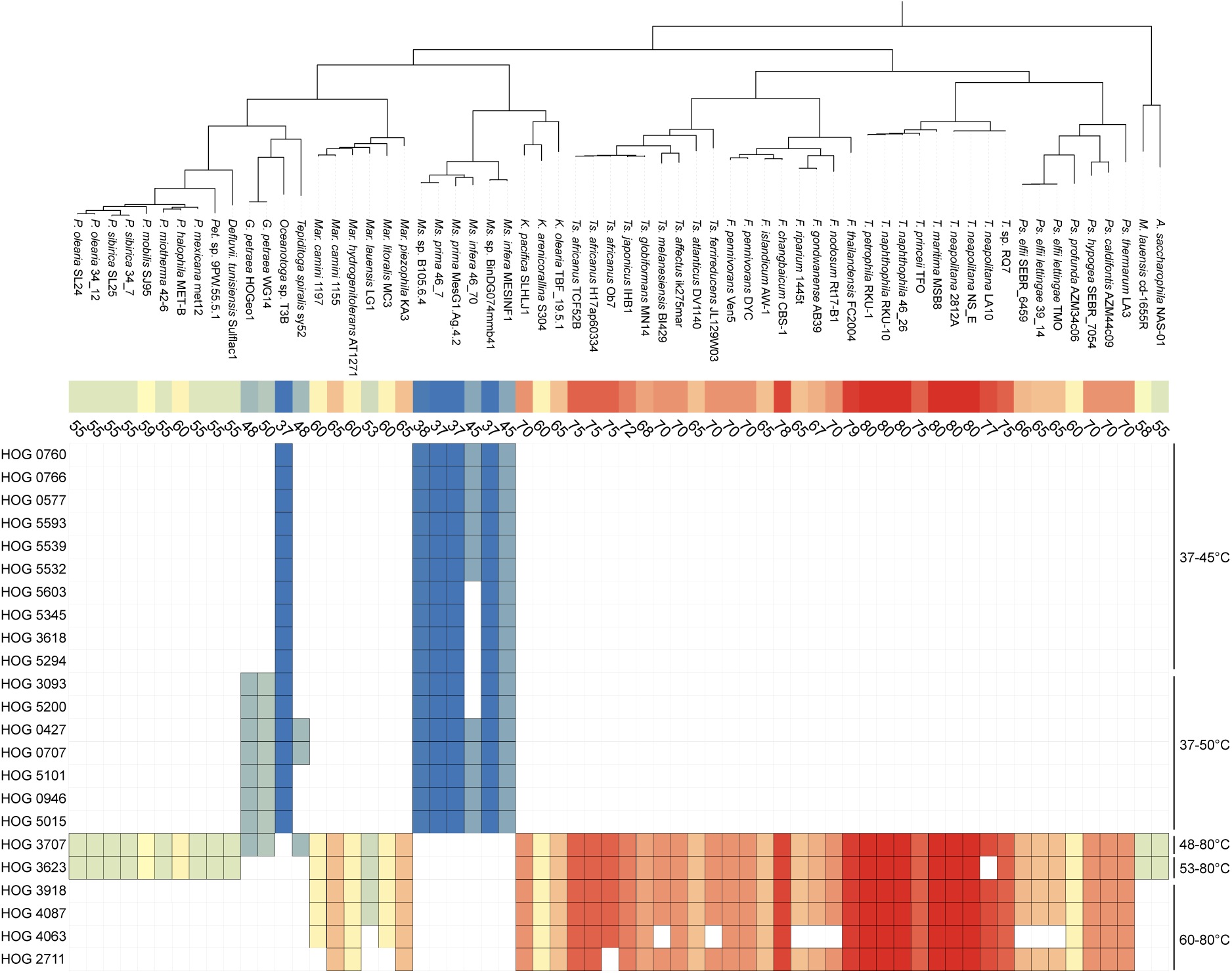
The presence and absence of 23 gene families with the strongest association with OGT across *Thermotogota*. Each row represents a gene family (HOG #) and each column corresponds to one of the 64 *Thermotogota* genomes, arranged according to the reference phylogeny (shown on top). Alongside each taxon name is its OGT in degrees Celsius and as a color square. For each gene family, presence of the gene in a taxon is indicated by a square colored by the taxon’s OGT. The families are grouped according to the temperature intervals with which the presence of the family is associated (shown on the right). Additional information about each gene family, including its functional annotation, is provided in **Table 1**.

**Table 1.**
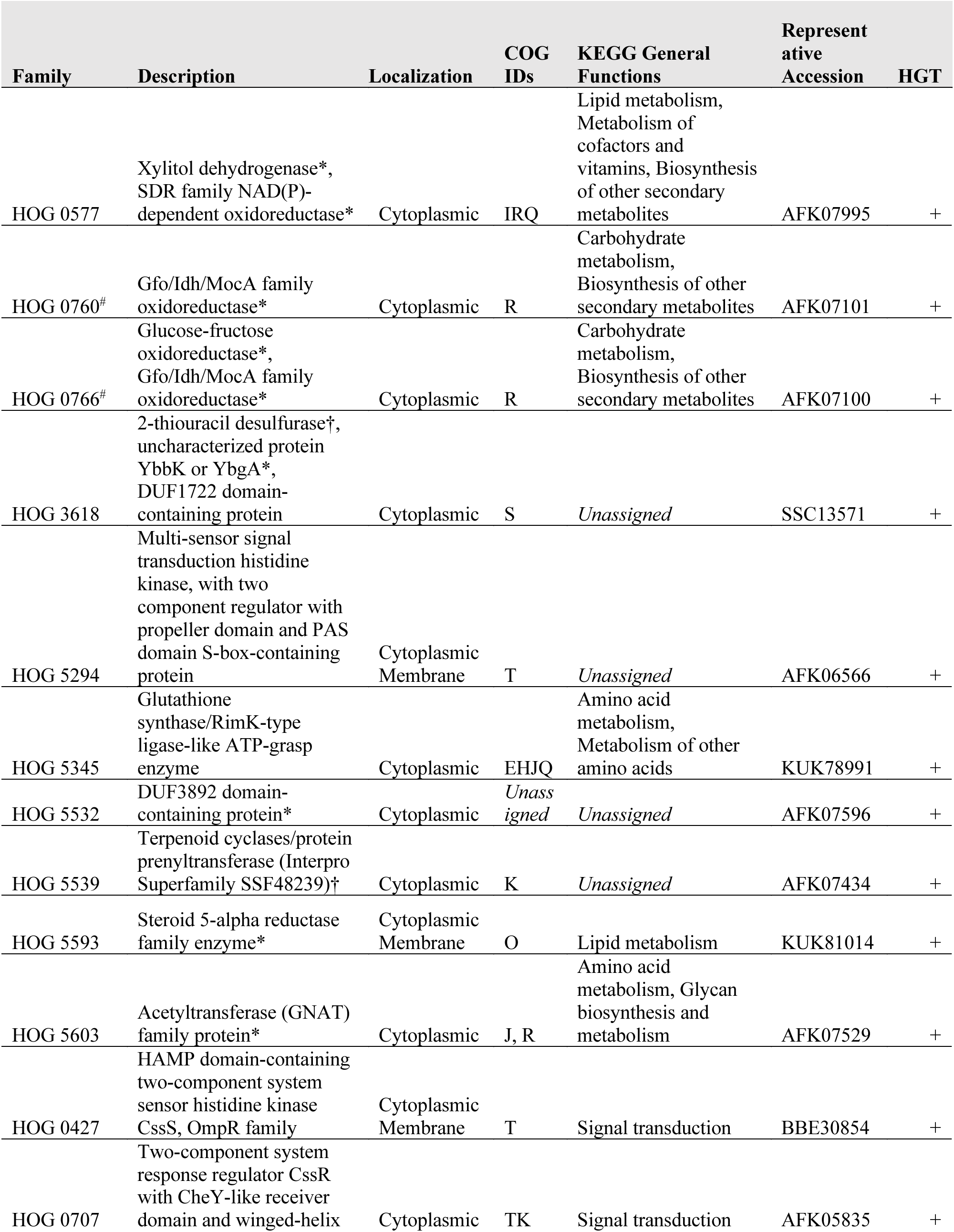
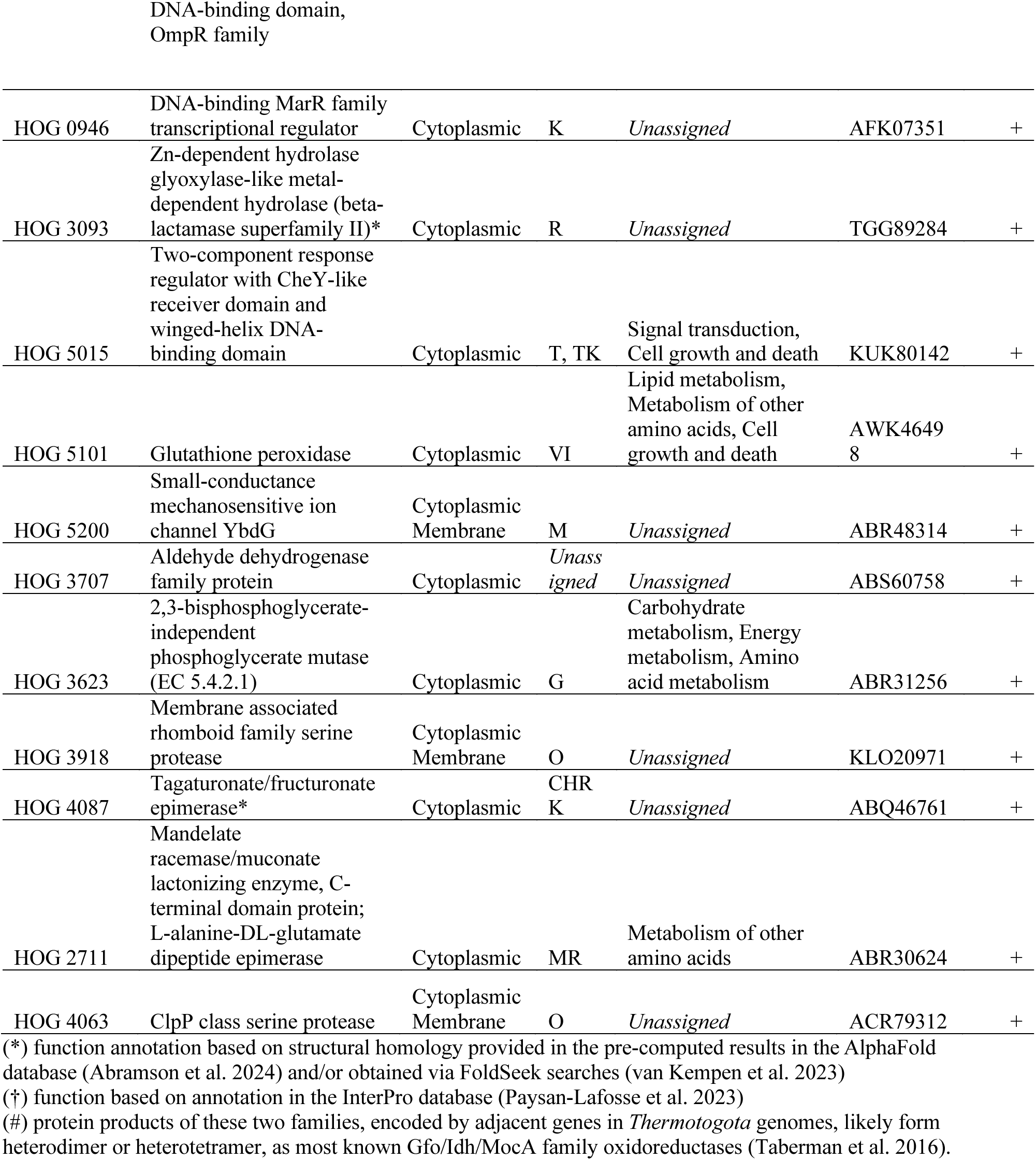
Functional annotations of the 23 gene families strongly correlated with specific OGTs.

The presence of 17 of the 23 gene families is associated with the lowest OGTs in the phylum: 10 families are found in most (and only) *Thermotogota* with OGT between 37 and 45°C, while the remaining 7 families are found in *Mesotoga, Oceanotoga*, *Geotoga,* and *Tepiditoga* species, which have OGTs between 37 and 50°C (**Figure 5**). Consistent with a hypothesis that acquisition of regulatory genes, signal transduction genes, secondary metabolite biosynthesis genes, and amino acid metabolism genes played an important role in the *Thermotogota*’s adaptation to mesophily (Zhaxybayeva et al. 2012), most of the 17 gene families fall into these functional categories (**Table 1**). Specifically, of the 10 families found in *Thermotogota* with OGT between 37 and 45°C, eight are involved in either carbohydrate metabolism (**HOG 0760** and **HOG 0766**), lipid metabolism (**HOG 0577** and **HOG 5593**), amino acid metabolism (**HOG 5345** and **HOG 5603**), nucleotide metabolism (**HOG3618)** or transcription and signaling (**HOG 5294**). **HOG 5539** is homologous to a family of terpenoid cyclases, which help cells regulate membrane fluidity and respond to common environmental stressors, including temperature (Avalos et al. 2022). Some terpenoids have also been shown to improve growth at low temperatures in *E. coli* and *Listeria* spp. (Avalos et al. 2022). The remaining gene family, **HOG5532**, is present in taxa outside of *Thermotogota* and encodes structurally conserved but functionally uncharacterized protein. Among 7 gene families that are found in *Mesotoga, Oceanotoga*, *Geotoga,* and *Tepiditoga* species, one is involved in lipid and amino acid metabolism as well as in defense against oxidative stress (**HOG 5101**), four are involved in transcription and signaling (**HOG 0427, HOG 0707, HOG 5015** and **HOG 0946**), one is involved in cell wall biogenesis (**HOG 5200)** and one is poorly characterized protein **(HOG 3093**). The *CssS hAMP* histidine kinase (**HOG 0427**) and the *CssR* DNA-binding response regulator (**HOG 0707**) encode the *CssRS* complex, which regulates heat shock proteases and is well-studied in *Bacillus* (Roncarati & Scarlato 2017; Noone et al. 2012; Sarvas et al. 2004; Darmon et al. 2002). It was proposed that *CssS* is a temperature sensor (Darmon et al. 2002). The two-component system responds to both heat and secretion stress in *Bacillus* and triggers the expression of chaperone-like proteases (Noone et al. 2012). The complex is important in managing the negative impacts of both extra-cytoplasmic misfolded proteins (Sarvas et al. 2004) and peptidoglycan stress (Dahal et al. 2023). Families **HOG 5015** and **HOG 0946** are transcription regulators from families known to be involved in response to environmental stimuli. A mechanosensitive ion channel (**HOG 5200**) could be triggered by temperature-induced stress, as bacterial mechanosensitive ion channels are activated by a stress to a cell membrane (Pliotas et al. 2015). These mechanosensitive proteins can alter gene expression (Harper et al. 2023), opening a possibility of **HOG 5200**’s involvement in temperature-sensitive gene regulation.

All 17 gene families are predicted to be horizontally exchanged, and 11 of them were likely independently acquired in different lineages of *Thermotogota* from other phyla (**Figure 6** and **Supplemental Figure S9**). Specifically, in 7 of the 10 families found in *Thermotogota* with OGT between 37 and 45°C, *Oceanotoga* str. T3B and *Mesotoga* spp. form separate groups, each closely related to taxa outside the *Thermotogota*. In 4 of the 7 gene families present in *Thermotogota* with OGTs between 37 and 50°C, *Geotoga*, *Oceanotoga,* and *Tepiditoga* form a distinct clade from *Mesotoga* species. Combined with the DTL-model inferences of horizontal transfer events in the evolutionary histories of these gene families, we conjecture that these genera have independently adapted to lower OGTs.

**Figure 6.**
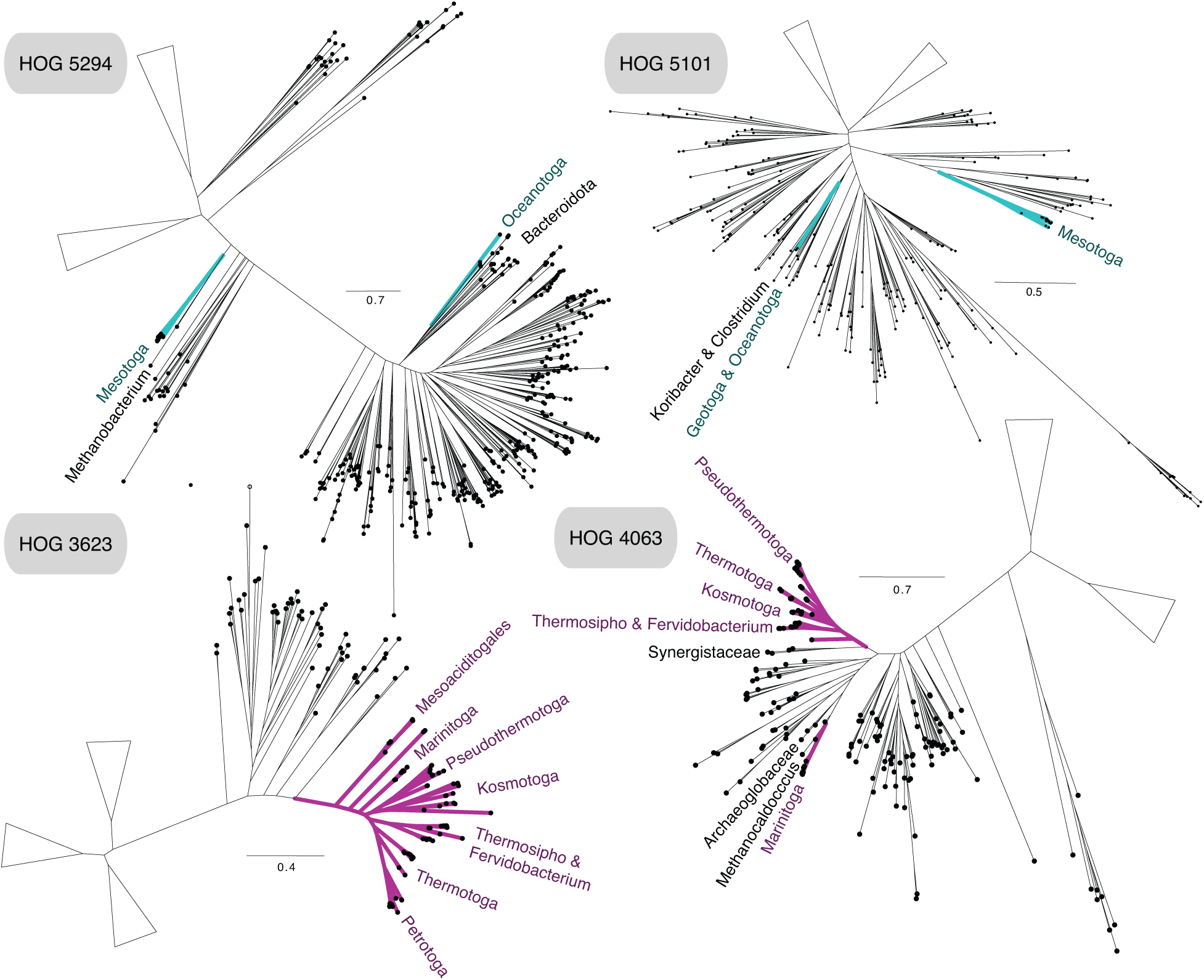
Expanded phylogenies of four gene families that illustrate various horizontal gene transfer scenarios. *Thermotogota* are highlighted in blue (for families present in taxa with OGTs <50°) or red (for families present in taxa with OGTs >50°). *Thermotogota* and their closest sister taxa are labeled. All phylogenies are unrooted, with some large groups of non-*Thermotogota* taxa collapsed into triangles. Complete phylogenies are available in the *FigShare* repository.

Among the remaining 6 of the 23 gene families, an uncharacterized member of the aldehyde dehydrogenase superfamily (**HOG 3707**) is present in all thermophilic *Thermotogota* (OGT of at least 48°C) and absent in all mesophilic *Thermotogota*. On the expanded phylogeny all *Thermotogota* group together, although the DTL model infers one transfer event between *Petrotoga* species. The phylogeny of the gene family, which is largely congruent with the reference phylogeny, indicates that the gene very likely was lost from *Mesotoga* spp. and *Oceanotoga* str. T3B (**Supplemental Figure S9**).

The gene family encoding 2,3-bisphosphoglycerate-independent phosphoglycerate mutase (**HOG 3623**), one of the central enzymes in the glycolysis pathway, is found exclusively in *Thermotogota* with OGT of at least 53°C. Notably, the gene family encoding an alternative 2,3-diphosphoglycerate-dependent phosphoglycerate mutase (**HOG 5300**) is associated primarily with OGT of 37-45°C (**Supplemental Table S3**). This enzyme has been implicated in heat sensitivity of a spirochaete (Benoit et al. 2001). Interestingly, *A. saccharophila* (OGT 55°C) has both **HOG 3623** and **HOG 5300**, suggesting that it may utilize different versions of this enzyme depending on environmental conditions. On the **HOG 3623**’s expanded phylogeny all *Thermotogota* group together, however the relationships among *Thermotogota* genera indicates a history of gene transfer within the phylum (**Figure 6**), which is supported by the 4 transfers inferred by the DTL model.

Finally, four genes are found predominantly in *Thermotogota* with OGT above 60°C: a tagaturonate/fructuronate epimerase (**HOG 4087**), a membrane-associated rhomboid family serine protease (**HOG 3918**), a *ClpP*-family serine protease (**HOG 4063**), and L-alanine-DL-glutamate epimerase (**HOG 2711**). While annotated as “radical SAM domain-containing protein”, **HOG 4087** is structurally homologous to tagaturonate/fructuronate epimerase (UxaE), thus representing a temperature-specific enzyme likely involved in carbohydrate metabolism.

The *ClpP*-family proteases (**HOG 4063**) assist in the unfolding and denaturing of proteins and are important in the management of cell stress (Msadek et al. 1998; Knudsen et al. 2014; Kwon et al. 2003). A *ClpP* gene is required by *Bacillus subtilis* for growth at higher-than-optimal temperatures (Msadek et al. 1998), promotes growth in *Streptococcus pneumoniae* under heat shock (Kwon et al. 2003), is upregulated in *E. coli* in response to heat stress (Pysz et al. 2004), and is important for cold tolerance in *Salmonellae* (Knudsen et al. 2014). Furthermore, a *ClpP* is down-regulated during heat shock in *T. maritima* (Pysz et al. 2004) and homologs from this gene family are upregulated at non-optimal temperatures in *Kosmotoga olearia* (Pollo et al. 2017), indicating that in *Thermotogota* it may help thermophiles cope with temperature stress. *ClpP* family proteases is a large family with multiple paralogs found in most bacteria: the *Thermotogota* pangenome includes 17 additional gene families annotated as *ClpP*. Yet, in BLASTP searches with the **HOG 4063** gene family as queries, other *Thermotogota ClpP*-family genes are not retrieved. *Kosmotoga* and *Marinitoga* species likely gained the **HOG 4063** family via transfer events (**Figure 6**). Notably, in the expanded phylogeny, *Marinitoga* spp. are sister taxa to hyperthermophilic archaeon *Methanocaldococcus infernus* (OGT of 85°C) (**Figure 6**).

Rhomboid family proteases (**HOG 3918**) are not well characterized in prokaryotes, but a recent study showed that in *Corynebacterium* spp. rhomboid protease genes are upregulated in response to heat stress and hypothesized that these proteases play a role in modification of the cell wall (Luenenschloss et al. 2022). Both the DTL model and the expanded gene family phylogeny suggest that species within the *Thermotogales* order inherited their **HOG 3918** homologs vertically, while *Kosmotoga* and *Marinitoga* species gained the family via transfer events (**Supplemental Figure S9**). Notably, a group of *Marinitoga* species is closely related to three *Bacillota* species, *Carboxydothermus hydrogenoformans*, *Caldicellulosiruptor changbaiensis* and *Caldicellulosiruptor kronotskyensis* (OGTs 70-75°C).

The L-alanine-DL-glutamate epimerases (**HOG 2711**) play a role in cell wall biogenesis and therefore may contribute to temperature tolerance by changing cell wall stability. This gene family includes a homolog from *Kosmotoga olearia* that is differentially transcribed at different temperatures (Pollo et al. 2017). While all *Thermotogota* form a group separate from other taxa on the expanded phylogeny, *Marinitoga* spp. may have gained this gene family from *Thermosipho* spp. (**Supplemental Figure S9**).

Taken together, the phylogenies of the latter 3 gene families raise a possibility that the *Marinitoga* spp.’s adaptation to higher OGTs in comparison to the rest of *Petrotogales* may be driven by gains of genes from thermophilic taxa.

## DISCUSSION

Our three independent estimates of ancestral OGT indicate that the LCTA was a thermophile, rather than the hyperthermophile it is previously presumed to be (Zhaxybayeva et al. 2009; Butzin et al. 2013; Green et al. 2013). As *Thermotogota* diversified and occupied different ecological settings, multiple independent increases and decreases of OGT led to presence of thermophiles, mesophiles and hyperthermophiles in the phylum. Our findings are consistent with an earlier analysis of 33 *Thermotogota* (Dahle et al. 2011).

We found that changes in OGT are associated with numerous gains and losses of genes throughout *Thermotogota*’s evolutionary history. We identified 597 gene families, whose presence and absence are associated with *Thermotogota*’s growth at different temperatures, and for 68 of these gene families the correlation is particularly convincing. Our findings underscore that adaptation to growth at different temperature requires products of multiple genes.

HGT has been documented as a driving force for microbial adaptations, from promoting diversification of microbes in hyperthermophilic environments (van Wolferen et al. 2013) to facilitating the spread of genes addressing a variety of environmental pressures (Arnold et al. 2022), such as antibiotic resistance (Woods et al. 2020). The majority of the 68 OGT-associated gene families have complex phylogenetic relationships that are not congruent with evolutionary history expected by vertical inheritance and speciation. More than 70% of the 68 gene families were not present in the LCTA and likely have been gained via HGT from outside of the phylum. Many of the potential donors of genes are *Bacillota* and archaea. This is consistent with the earlier studies that suggested a substantial exchange of genes between *Thermotogota* and archaea (Pollo et al. 2015), and between *Thermotogota* and *Bacillota* (Zhaxybayeva et al. 2009). Most of the genes inferred to be present in LCTA also experienced multiple gene transfer events within the phylum. Taken together, our analyses showed that HGT plays a significant role in evolution of OGT in *Thermotogota*, and there is likely a pool of temperature-specific genes in a microbial community, acquisition of which allows *Thermotogota* (and likely other bacteria and archaea in the community) to adapt to new growth temperature conditions. In support of our findings, reverse gyrase gene (which is necessary for life above 90°C (Lipscomb et al. 2017) and possibly for hyperthermophily in general) was shown to be acquired horizontally multiple times (Catchpole & Forterre 2019).

The products of the 68 OGT-associated genes contribute to diverse metabolic functions (amino acid metabolism, carbohydrate metabolism, lipid metabolism and energy production), signal transduction and transcription regulation. Some of the genes could, of course, be spuriously correlated with OGT and instead contributing to other phenotypic traits. Yet, these functional assignments are consistent with existing evidence for cellular processes impacted by growth temperature. For example, in *Thermotogota*, the species *Kosmotoga olearia* and *Thermotoga maritima* shift their carbohydrate metabolisms when temperature is not at optimum (Wang et al. 2012; Pollo et al. 2017). In *Thermosipho africanus,* a large set of carbohydrate metabolism genes has been hypothesized to contribute to their habitat difference from other *Thermosipho* spp. (Haverkamp et al. 2017). Such metabolic adaptations are not limited to *Thermotogota*: for example, *E. coli* adapted their metabolic networks to new environments by gaining genes involved in the transport and catalysis of external nutrients (Pál et al. 2005), and in *Thermus filiformis*, carbohydrate metabolism changed as a result of experimental thermoadaptation (Mandelli et al. 2017). Lipid membrane integrity, structure, and function are highly impacted by temperature (Siliakus et al. 2017). Membrane transport systems play a major role in cell efficiency and adaptation because they facilitate a cell’s exchange with its environment (Rees et al. 2009). Temperature-adapted transport systems can reduce metabolic costs and make environmentally-specific substrates available (Rees et al. 2009). Finally, adjusting cellular processes based on environmental cues is generally critical in adaptive responses (Stock et al. 1989; López-Maury et al. 2008), and thus could explain the presence of temperature/environment sensors and transcription factors among the 68 genes.

Occurrence of a few either uncharacterized or general-function-prediction-only genes in our results, even after searches for structural homologs, suggests that some proteins that contribute to ability to grow at a specific temperature remain unknown. Further investigations of these genes’ functions are needed to explain their connections with OGT.

Notably, the identified 68 genes are associated with a variety of overlapping temperature intervals. While traditionally the organisms are classified into psychrophiles, mesophiles, thermophiles and hyperthermophiles using arbitrarily-set temperature cutoffs, it is likely that multiple, gradual adaptations are needed across temperature spectrum rather than at discrete temperature thresholds.

## METHODS

### Retrieval of *Thermotogota* genomes, assessing completion, and assignment of optimal growth temperature

A dataset of 157 non-redundant *Thermotogota* genomes and metagenome-assembled genomes (MAGs) and 15 outgroup bacterial genomes, which we previously curated (Farrell et al. 2023), was updated with 2 additional *Thermotogota* genomes (*Thermosipho ferrireducens* JL129W03 and *Fervidobacterium riparium* 1445t) obtained from the NCBI’s Assembly database in October 2022 (**Supplemental Tables S1** and **S6**). The dataset includes genomes from 48 validly described *Thermotogota* species. Genomes were assigned taxonomic ranks using the Genome Taxonomy Database (GTDB) (Chaumeil et al. 2020). MAGs and draft genomes were checked for completion and contamination using CheckM (Parks et al. 2015).

The OGTs of *Thermotogota* isolates was obtained from the literature (see **Supplemental Table S1** for sources). To assign OGT to as many genomes as possible, the OGT of a species was attributed to genomes of all strains of the same species, unless a strain’s specific OGT was known. This procedure resulted in 64 *Thermotogota* genomes with an associated OGT (**Supplemental Table S1**), with at least one representative genome from each genus of *Thermotogota*.

### Reconstruction of phylogenies from 50 concatenated ribosomal proteins

Amino acid sequences of translated ribosomal protein genes were identified by BLASTP v.2.6.0 (Altschul et al. 1990) searches of 50 *Thermotoga maritima* ribosomal proteins against the database of the above-described 159 *Thermotogota* (**Supplemental Table S1** and **S6**) and 15 outgroup (**Supplemental Table S7**) genomes. Matches were kept if they were annotated as ribosomal proteins, had an e-value <0.0001, and had the subject length between 60 and 140% of the query length. For each of the 50 ribosomal proteins, the retrieved homologs were aligned in MAFFT-linsi v7.305b (Katoh & Standley 2013).

The 50 alignments were concatenated into one alignment (8,092 amino acid residues long) using in-house scripts; gaps were inserted into the alignment if a genome did not have a detected ribosomal protein. The phylogenetic tree was reconstructed in IQ-TREE v1.6.12 (Nguyen et al. 2015) using the concatenated alignment and the LG+F+R6 substitution model selected by built-in ModelFinder (Kalyaanamoorthy et al. 2017). To assess accuracy of the reconstructed tree, bootstrap analysis of 100 pseudo-samples was carried out. The phylogeny was rooted using the outgroup taxa and used to detect gene families (see **“Identification of gene families”** section below).

For the second phylogeny, used as the reference phylogeny throughout the paper, the concatenated alignment of only 64 *Thermotogota* strains with assigned OGTs and the 15 outgroup genomes, was created (7,790 amino acid residues long). The phylogenetic tree was reconstructed in IQ-TREE v1.6.12 (Nguyen et al. 2015) under the LG+F+R5 substitution model selected by built-in ModelFinder (Kalyaanamoorthy et al. 2017).

### Testing models to describe evolution of OGT

The rooted reference phylogeny was converted to an ultrametric tree using the *chronoMPL* function of the “ape” R package (Paradis & Schliep 2019; Britton et al. 2002). Pagel’s lambda was estimated using the *phylosig* function of the “phytools” R package (Revell 2012). A Brownian Motion model (Felsenstein 1973), an Early Burst model (Blomberg et al. 2003), and an Ornstein-Uhlenbeck model (Hansen 1997) were fit to the ultrametric tree using the *fitContinuous* function from the “geiger” R package (Pennell et al. 2014). The fit of the models to the data was compared using weighted AIC values (Harmon et al. 2010).

### Reconstruction of OGT as a continuous trait along the *Thermotogota* phylogeny

Reconstruction of ancestral OGT was performed using BayesTraits v3.0.2 (Pagel et al. 2004) on the reference phylogeny under a random walk. The model was run using Markov-Chain Monte Carlo (MCMC) with 500 stones (2,000 iterations each), 100,000,000 iterations, and a prior for alpha set at (−100;100) range. A burn-in was set to 20,010,000 trees after an inspection of log-likelihood plots for convergence. The OGT of the LCTA was calculated as the mean OGT value for common ancestral node of all *Thermotogota* among the remaining trees, and the credible interval was calculated as 95% of posterior distribution OGT values (Huelsenbeck et al. 2002; Huelsenbeck & Bollback 2001).

### Reconstruction of OGT from the GC content of the 16S rRNA gene

A phylogeny of 16S rRNA genes was reconstructed using taxa matching the reference phylogeny. The sequences were retrieved by querying *Thermotoga maritima*’s 16S rRNA gene (NR_102775.2) against each of the 64 *Thermotogota* genomes using BLASTN v.2.6.0 (Altschul et al. 1990) with default settings (E-value<10). For each genome, the match >1,000 nt long and with the highest percent identity was extracted using in-house scripts. For 9 genomes with no matches to the query, sequences to represent the strain were searched for in the NCBI’s Nucleotide database using text searches for species and strain names (accessed in December 2023). As a result, 6 of the 9 genomes were represented by a 16S rRNA gene either from the same strain or from the alternate strain of the same species; the other 3 genomes remained unrepresented. This resulted in 61 16S rRNA gene sequences matched to the 64 genomes in the reference phylogeny (**Supplemental Table S1**). Fifteen 16S rRNA gene sequences for the outgroup taxa (**Supplemental Table S7**) were retrieved from the NCBI’s Nucleotide database. The 16S rRNA gene sequences were aligned with SILVA Incremental Aligner v1.2.11 (SINA) (Pruesse et al. 2012). The maximum likelihood (ML) tree, nonparametric bootstrap samples, and ancestral states for all nodes of the tree were reconstructed using IQ-TREE (Nguyen et al. 2015) under the GTR+F+R4 model as selected by ModelFinder (Kalyaanamoorthy et al. 2017). The resulting phylogeny has a previously identified artefact in *Thermotogota* 16S rRNA gene trees (Farrell et al. 2023), where the *Kosmotogaceae* family groups closest to the *Mesoaciditogales* order instead of being a sister clade to the *Petrotogaceae* family (**Supplemental Figure S3**).

To eliminate the artifact, as well as to include 12 described *Thermotogota* species that are not represented in our reference phylogeny due to lack of sequenced genomes, an alternate dataset of 16S rRNA genes from 60 *Thermotogota* species was assembled. Only one representative 16S rRNA gene sequence for each species with an assigned OGT was included, and the sequences from the type strains were chosen over other available strains where possible (**Supplemental Table S8** and **Supplemental Figure S2**). While the taxa in this 16S rRNA gene dataset and the ribosomal proteins dataset are not 1:1, all genera are represented in both datasets (**Supplemental Table S8**). Eight outgroup taxa were used (**Supplemental Table S9**). The sequences were extracted from NCBI’s Nucleotide database and aligned with SINA v1.2.11 (Pruesse et al. 2012). The ML tree, nonparametric bootstrap samples, and ancestral states for all nodes of the tree were reconstructed using IQ-TREE (Nguyen et al. 2015) under the homogeneous, GTR+F+I+G4 model as selected by ModelFinder (Kalyaanamoorthy et al. 2017).

Ancestral reconstruction under non-homogeneous model was performed on both “64 *Thermotogota* strains” and “60 *Thermotogota* species” datasets using the “baseml” program of PAML v.4.9d (Yang 2007), with the above-described SINA alignments of extant sequences and the homogenous IQ-TREE phylogenies rooted with the outgroup as a guide. The “nhomo” parameter was set to 4 to model different sets of frequency parameters for the root and each branch on the tree, and a single rate was used across sequence sites (no genes or partitions).

The following analyses were performed for the ancestral reconstructions both under homogeneous and non-homogeneous models. For each tree node, each site of the ancestral sequence at the node was assigned the nucleotide with the highest posterior probability. The extant and reconstructed ancestral sequences were aligned together with a bacterial 16S rRNA scaffold from the Ribosomal Database Project (Cole et al. 2014) using “cmalign” program from Infernal v1.1.2 (Nawrocki & Eddy 2013). Stem regions from the resulting structural alignment were extracted using in-house scripts and their GC-content was calculated for both ancestral nodes and extant taxa. Linear regression was performed between the stem GC-content of extant taxa (**Supplemental Table S8**) and their OGTs using Python package *scikit-learn* 0.23.2 (Pedregosa et al. 2011). The fitted linear model was used to predict ancestral OGT based on the stem GC-content of ancestral nodes. The reliability of regression was tested by the phylogenetic independent contrasts analysis (Felsenstein 1985) as implemented in the *ape* package in R (Paradis & Schliep 2019).

### Identification of gene families

Gene families in 174 genomes (159 *Thermotogota* genomes and MAGs and 15 outgroup taxa; **Supplemental Tables S1, S6** and **S7**) were detected using OrthoFinder v.2.5.1 (Emms & Kelly 2019). Within OrthoFinder, BLASTP v.2.6.0 was used for the sequence search, MAFFT v.7.305b for sequence alignment, and the concatenated ribosomal proteins phylogeny as the reference phylogeny. The gene families were defined as the hierarchical orthogroups (HOGs) for “N0”.

A total of 13,121 gene families (excluding ORFans) were detected in the 174 genomes. Of those families, 6,264 gene families are present in the subset of 64 *Thermotogota* genomes with assigned OGT.

### Identification of genes and gene families encoding transmembrane proteins

Transmembrane regions were predicted for amino acid sequences of all protein-coding genes in the genomes of the 64 *Thermotogota* with assigned OGT using Phobius v.1.01 (Käll et al. 2005). Genes that had a TM score ≥3 were designated as transmembrane proteins. A gene family was designated as a transmembrane protein family, if it comprised >50% genes predicted to encode transmembrane proteins, and as a non-transmembrane family otherwise.

### Ancestral proteome reconstruction

For each non-transmembrane family present in ≥75% of the 64 *Thermotogota* genomes with assigned OGT, amino acid sequences were aligned using MAFFT-auto v7.305b (Katoh & Standley 2013) and ML tree was reconstructed using IQ-TREE v.1.6.12 (Nguyen et al. 2015). Trees were rooted with the outgroup taxa using in-house scripts. Gene families whose trees could not be rooted because either the family did not contain any sequences from outgroup taxa or if less than 3 outgroup taxa grouped together, were excluded. To ensure the inclusion of the LCTA node, gene families that did not have genes from at least one species from each of the three *Thermotogota* orders (*Thermotogales, Petrotogales* and *Mesoaciditogales*) were also excluded.

For the resulting 556 gene families, the LCTA node was identified as the common ancestral node for the 64 *Thermotogota* taxa. Because *Thermotogota* have undergone extensive HGT throughout their history (Nesbø et al. 2009; Zhaxybayeva et al. 2009) and their gene families are known to have complicated phylogenies (Farrell et al. 2023), it was also checked that the *Mesoaciditogales* diverged from the LCTA node before the *Thermotogales* and *Petrotogales* orders split. For 294 of 556 gene family trees that met this criterion, ancestral states for all nodes were reconstructed using IQ-TREE v.1.6.12 (Nguyen et al. 2015). For ancestral sequences within each family, the proportion of IVYWRELGKP amino acids was calculated by summing the probabilities of these amino acids in each site and dividing the sum by the length of the reconstructed protein.

### Correlation between amino acid content and OGT of extant and ancestral *Thermotogota*

Median IVYWRELGKP was calculated for all non-transmembrane proteins encoded in the genomes of the 64 *Thermotogota* with an assigned OGT using in-house scripts. Linear regression was performed to establish a correlation between IVYWRELGKP and OGT. The median value of the 294 reconstructed ancestral protein sequences was used to predict the LCTA’s OGT using the regression model. A 95% confidence interval was used when computing the t-score for calculating the margin of error for the predicted OGT of the LCTA.

### Modeling gains, losses, and expansions of gene families using the COUNT program

A matrix of gene counts per family per genome was created from 4,746 gene families found in more than 3 of the 64 *Thermotogota*. The gene count matrix and reference phylogeny were used as input for the COUNT v.10.04 (Csűös 2010). All combinations of models with family-specific distribution parameters ranging from 1 to 3 were compared using likelihood ratios. The chosen ML model used a constant rate for all family-specific rates, with maximum paralogs set to 300. Gene family presence, gains, losses, expansions, and reductions were considered “events” when their posterior probabilities were ≥60%. Genomes with low completion can return inflated loss values at their leaves, therefore values on terminal branches leading to genomes with <90% completion were removed from loss correlation analyses. Trends between gene family dynamics and the estimated OGT shifts from the random walk model were examined using in-house scripts.

### Testing Association of Gene Presence/Absence with OGT

Presence/absence of genes that significantly associate with OGT were identified using the Fixed Effects model of Pyseer v.1.3.9 (Lees et al. 2018). The following data were used as input into Pyseer: a set of 3,604 unique presence-absence patterns that was extracted from a subset of 6,081 gene families found in at least 2 of the 64 *Thermotogota*, OGT values as a continuous trait, and a distance matrix generated from the reference phylogeny of 64 *Thermotogota* with assigned OGTs. Using the scree plot as a guide (**Supplemental Figure S10**), the maximum dimension for multi-dimensional scaling was set to 8, representing the largest value that still produced significant associations. The p-value of 1.42E-05 was selected as a threshold for significance, using a script in the Pyseer package. To look for the strongest associations with OGT, the phyletic patterns associated with OGT were curated as follows. Families that were either present only in taxa from a single genus or family, as well as families that were absent only in taxa from a single genus or family were filtered out. Remaining families were further screened to select families that have the best correlations with OGT in three intervals: 37-45°C, 48-59°C, and 60-80°C. Due to different number of *Thermotogota* in each OGT range, different criteria were used for each interval. To select families best correlated with OGT of 37-45°C, 60% of the taxa in a family were required to have OGTs of 37-45°C and contain fewer than 5 taxa with OGTs of 60-80°C. To select families best correlated with OGT of 48-59°C, at least 50% of the taxa in a family were required to have OGTs of 48-59°C. To select families best correlated with OGT of 60-80°C, 60% of the taxa were required to have OGTs of 60-80°C, contain no taxa with OGTs of 37-48°C, and required to be detected in at least 25% of *Thermotogota* with OGTs of 60-80°C.

### Duplication-Transfer-Loss modeling

Each gene family was filtered to include amino acid sequences only from 64 *Thermotogota* with assigned OGTs. For each family, the sequences were aligned using MAFFT-auto v7.305b (Katoh & Standley 2013) and phylogeny reconstructed using IQ-TREE v1.6.12 (Nguyen et al. 2015) under the best-fitting model identified via ModelFinder (Kalyaanamoorthy et al. 2017). The origins, duplications, transfers, and losses of the OGT-associated gene families were inferred using a species-tree-aware method, as implemented in GeneRax v2.1.2 (Morel et al. 2020). Rooted reference phylogeny of the 64 *Thermotogota* and 15 outgroup taxa was provided as the “species tree”. For each gene family, sequence alignment was used as data and its unrooted phylogeny as a starting tree. The best-fit model chosen during the phylogenetic reconstruction of the starting tree was used as the substitution model. In the two cases where the suggested model (mtInv+F+I+G4) was not part of the GeneRax suite, the default LG+G4 model was used. The “Undated DTL” model, per-family rates, and the SPR tree search strategy options were selected.

### Functional annotation of OGT-associated gene families

Cellular localization of proteins encoded by a gene family was predicted via the DeepLocPro v1.0 (Moreno et al. 2024) web interface (accessed March 24, 2024), using as input one amino acid sequence of the median length as a gene family’s representative. Significant enrichment and depletion of cellular localization in the set of 68 genes families was assessed using a two-sided Fisher’s Exact Test, with a p-value threshold of ≤0.05.

Protein-coding genes from the 64 *Thermotogota* genomes were assigned COG categories by searching the amino acid sequences of the genes against the NCBI COG database (2020 release; downloaded in March 2021) (Galperin et al. 2021) using BLASTP v.2.6.0+ (Altschul et al. 1990) with e-value of 1e-10 and maximum target sequences set to one. COG categories of a family were defined as the set of COG categories that were assigned to the genes of that family.

Additionally, protein-coding genes from the 64 *Thermotogota* genomes were functionally annotated with KEGG terms using eggNOG-Mapper’s “emapper” v.2.1.3 (Huerta-Cepas et al. 2017) and Diamond v.2.0.9.147 search (Buchfink et al. 2021) of the eggNOG database (downloaded in June 2021). KEGG terms for each gene family were summarized using the KEGG Mapper web interface (accessed February 2024) (Kanehisa & Goto 2000; Kanehisa et al. 2022). KEGG Pathway matches specific to eukaryotes or human diseases were excluded.

Twenty-eight of the 68 gene families remained uncharacterized because they fell into at least one category: (1) assigned COG categories R or S, (2) not assigned to any COG category, or (3) their annotations included terms “hypothetical”, “putative”, or “domain of unknown function” (DUF). To improve their functional annotations, structural homology searches were carried out using the amino acid sequence with the median length as a query.

First, the query proteins were submitted to the AlphaFold (Abramson et al. 2024) web interface (accessed in July 2024). The functional annotation of the top pre-computed structural homolog in AlphaFold’s “Structure Similarity Cluster” was recorded. For 14 gene families, these annotations were either uncharacterized or contained only DUFs. Therefore, additional structural homology searches were performed as follows. Models of tertiary structures of the query proteins were extracted from the ESM Atlas v. 2023_02 using FoldSequence tool of the ESM Atlas API (Lin et al. 2023). The retrieved models were searched using FoldSeek (van Kempen et al. 2023) against the AlphaFold/UniProt50 database “AFDB50” (Varadi et al. 2022; The UniProt Consortium 2015) by submitting the queries via the FoldSeek API (van Kempen et al. 2023).

Matches with the e-values ≤ 0.001 and probabilities ≥ 0.8 were retained. The improved functional annotation of the query was selected manually from the results by assessing the overall top match, the best match outside the *Thermotogota* phylum, and the top match with a functional annotation that was not “hypothetical”, “DUF”, or “putative” (**Supplemental Table S10**).

For genes with no improved functional annotations from the above-described structural homology searches, the InterPro database (accessed via https://www.ebi.ac.uk/interpro/ in August 2024) (Paysan-Lafosse et al. 2023) searches and PHYRE2 searches (accessed in September 2024) (Kelley et al. 2015) were performed (**Supplemental Table S10**).

For each of the selected “best new annotation” match of a gene family, their Uniprot IDs were mapped to eggNOG, NCBI RefSeq, and KEGG using Uniprot’s id mapping tool (The UniProt Consortium 2015), and their amino acid sequences used as queries in searches against the NCBI’s Conserved Domain Database (Yang et al. 2020) and BlastKOALA (Kanehisa et al. 2016). COG or KEGG annotations obtained in these searches were assigned to a gene family (**Supplemental Table S11**).

### Reconstruction of expanded gene family phylogenies

For each of the 68 OGT-associated gene families, amino acid sequences of all proteins encoded by the gene family were used as queries in searches of the RefSeq database release 202 (O’Leary et al. 2016) using BLASTP in BLAST 2.6.0+ (Altschul et al. 1990), with the maximum number of target sequences set at 500 and all other settings as default. Matches with e-values ≤ 10^-6^ and ≥ 50% query and subject coverage were retained. For each gene family, the results of individual searches were combined, and duplicate matches were removed. Amino acid sequences of the retained matches were retrieved from NCBI using “efetch” v.15.6 from Entrez Direct (Kans 2024). To make tree building more efficient and make the large datasets more interpretable, sequences were clustered using MMSeqs2 v.12.113e3 (Steinegger & Söding 2017), using a maximum sequence identity threshold of 0.8 and all other parameters as default. Only one representative designated by MMSeq2 was retained for each cluster. If the *Thermotogota* sequences from a gene family were removed during clustering, they were added back.

For each expanded gene family, amino acid sequences were aligned using MAFFT v7.305 using the auto setting. ML trees were reconstructed in IQ-TREE v1.6.12 (Nguyen et al. 2015), using ModelFinder (Kalyaanamoorthy et al. 2017) to select the best substitution model and 1,000 ultrafast bootstraps to assess the tree topology support. The location of *Thermotogota* taxa in the tree was checked using in-house scripts that utilized the “Environment for Tree Exploration” v3 (ETE3) package (Huerta-Cepas et al. 2016) functions.

### Phylogenetic Tree Visualization

Phylogenetic trees were visualized an annotated using iTOL’s web platform v6 (Letunic & Bork 2021), ETE3 (Huerta-Cepas et al. 2016), FigTree v1.4.4 (Rachel Colquhoun et al. 2018), and ThirdKind (Penel et al. 2022).

## Data Availability

Most genomes and 16S rRNA sequences used for this research are publicly available via either NCBI Nucleotide (https://www.ncbi.nlm.nih.gov/nucleotide/), NCBI Assembly (https://www.ncbi.nlm.nih.gov/assembly) or the Joint Genome Institute Integrated Microbial Genomes (https://img.jgi.doe.gov/) databases (see **Supplemental Tables S1** and **S6** for accession numbers). From a few MAGs were shared with us but not yet publicly available, sequences of specific genes incorporated into gene families used in our analyses are available via a **FigShare** repository, as described below. The following data is available in a **FigShare** repository at DOI **10.6084/m9.figshare.26880394**: Amino acid sequences of proteins in each of the 13,121 gene families in FASTA format; list of 597 gene families significantly correlating with OGT; alignments and phylogenetic trees of the 68 selected gene families; alignments and phylogenetic trees of the 16S rRNA genes and concatenated ribosomal proteins; data associated with the ancestral reconstructions and ancestral OGT inferences from 16S rRNA gene and 294 gene families; gene presence/absence matrices used in the gain/loss and pangenome-wide association analyses; Python scripts used for data manipulation.

## Acknowledgements

We thank Eric Boyd (Montana State University), Håkon Dahle (University of Bergen), and Brett Baker (University of Texas in Austin) for sharing MAGs, which were used in gene family detection. The work was supported by Dartmouth Fellowship and Cramer funds to AAF, and Dartmouth Dean of Faculty funds to OZ.

## Notes

https://doi.org/10.6084/m9.figshare.26880394.v1

### Competing Interest Statement

The authors have declared no competing interest.

