## Supplemental Figures S1-S10 for "Bacterial growth temperature as a horizontally acquired polygenic trait"

**Supplemental Figure S1. Correlation between the OGT and GC content of the 16S rRNA stem regions in extant *Thermotogota*.** Each point represents one of the 60 described *Thermotogota* species with experimentally determined OGT. Trendline is shown as a red dashed line. **(a)** Linear correlation not corrected for phylogeny. **(b)** Linear correlation after correcting for phylogenetic relatedness. PIC, phylogenetically independent contrasts.

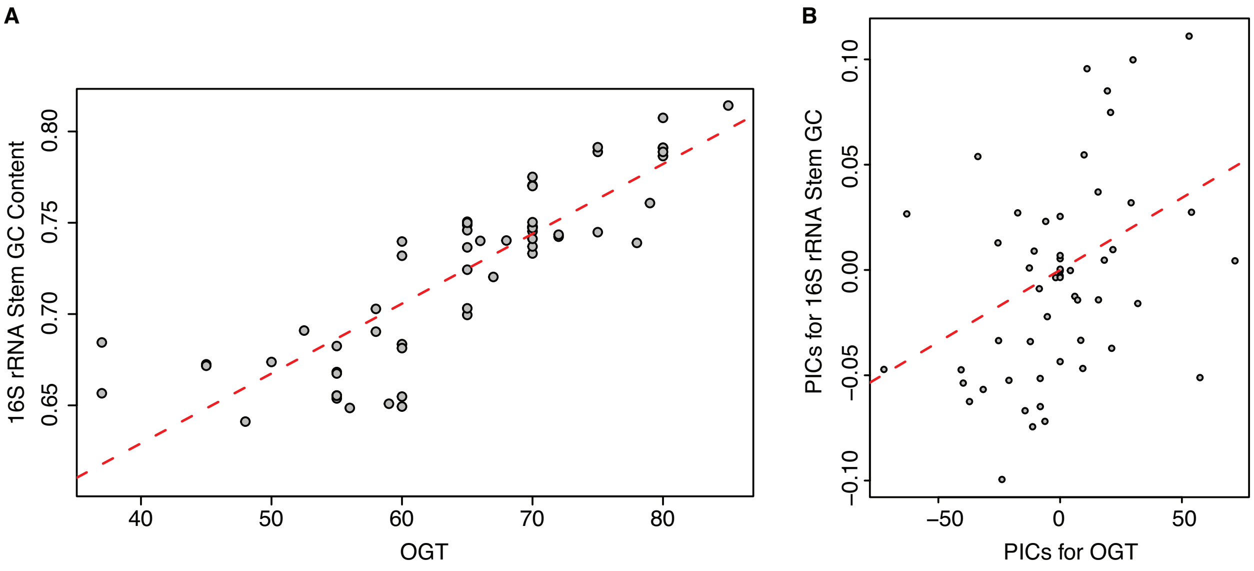

**Supplemental Figure S2. Predicted change in OGT from LCTA to 60 present-day *Thermotogota* species as inferred from GC content of reconstructed 16S rRNA stem regions.** OGT for each node was calculated using the linear regression shown in **Supplemental Figure S1** and mapped onto 16S rRNA gene phylogeny. Nodes are colored by either estimated (internal nodes) or extant (external nodes) OGT. OGT (in °C) is annotated in black and the GC content of the 16S rRNA stem regions (in %) is depicted in grey. The tree in Newick format that shows both bootstrap support values and OGT estimates with 95% confidence intervals is available in *FigShare* repository (see **Data Availability** section). Note that the topology of this 16S rRNA tree differs from the ribosomal protein phylogeny in **Figure 1:** *Thermosipho* spp. are sister taxa to *Thermotoga* and *Pseudothermotoga* spp. instead of *Fervidobacterium* spp., although with a low bootstrap support (79%). Scale bar, substitutions per site.

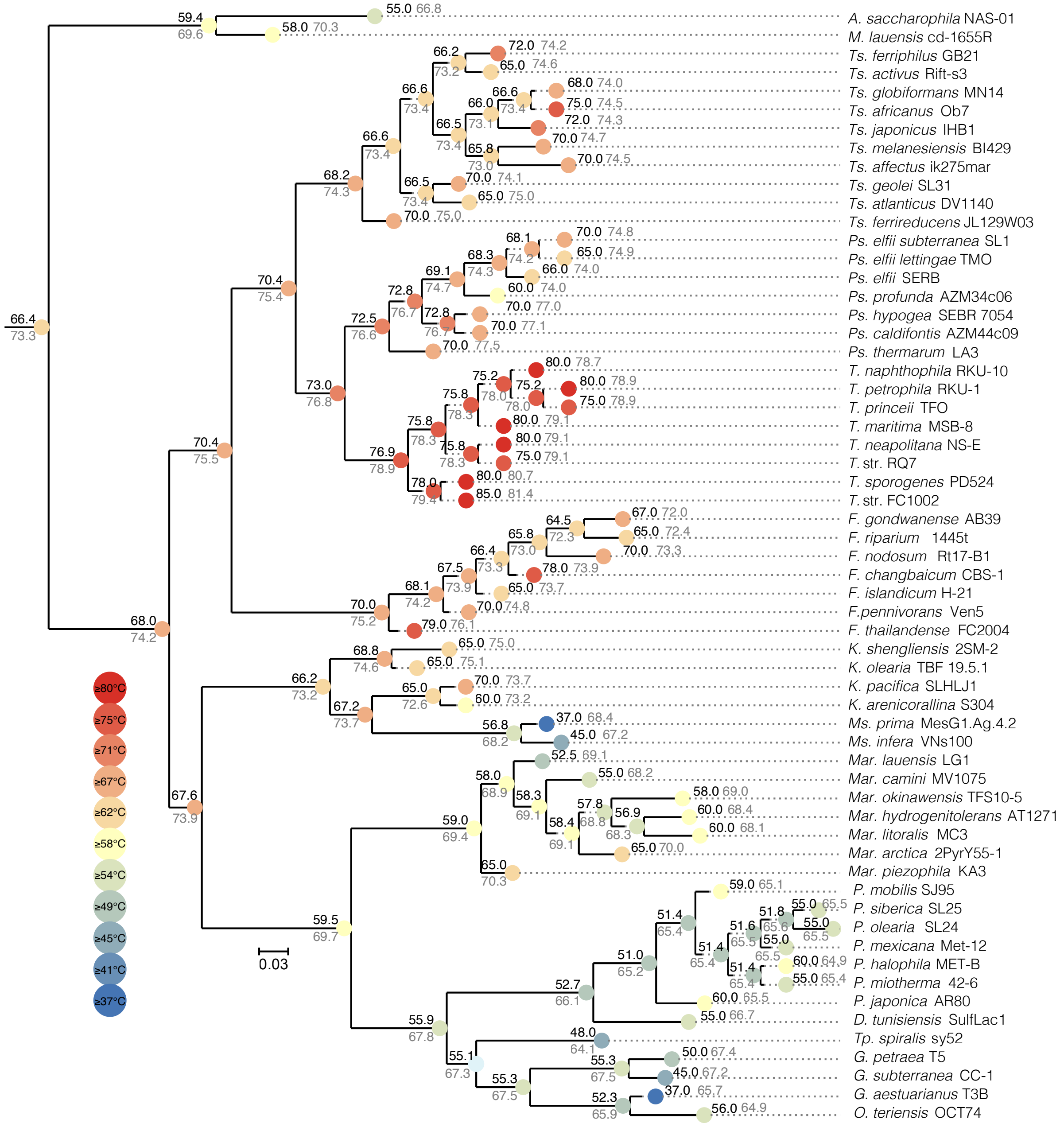

**Supplemental Figure S3. Predicted change in OGT from LCTA to 64 *Thermotogota* strains as inferred from GC content of reconstructed 16S rRNA stem regions.** OGT for each node obtained using the linear regression and mapped onto 16S rRNA gene phylogeny. Nodes are colored by either estimated (internal nodes) or extant (external nodes) OGT. OGT (in °C) is annotated above each branch and the GC content of the 16S rRNA stem regions (in %) is depicted below each branch. The tree in Newick format that shows bootstrap support values and OGT estimates with 95% confidence intervals is available in *FigShare* repository (see **Data Availability** section). Note that the topology of this 16S rRNA tree differs from the ribosomal protein phylogeny in **Figure 1:** *Thermosipho* spp. are sister taxa to *Thermotoga* and *Pseudothermotoga* spp. instead of *Fervidobacterium* spp., and *Kosmotogaceae* are a basally branching clade instead of being a sister group to *Petrotogaceae*. However, both discrepancies have low bootstrap support (59% and 41%, respectively). Scale bar, substitutions per site.

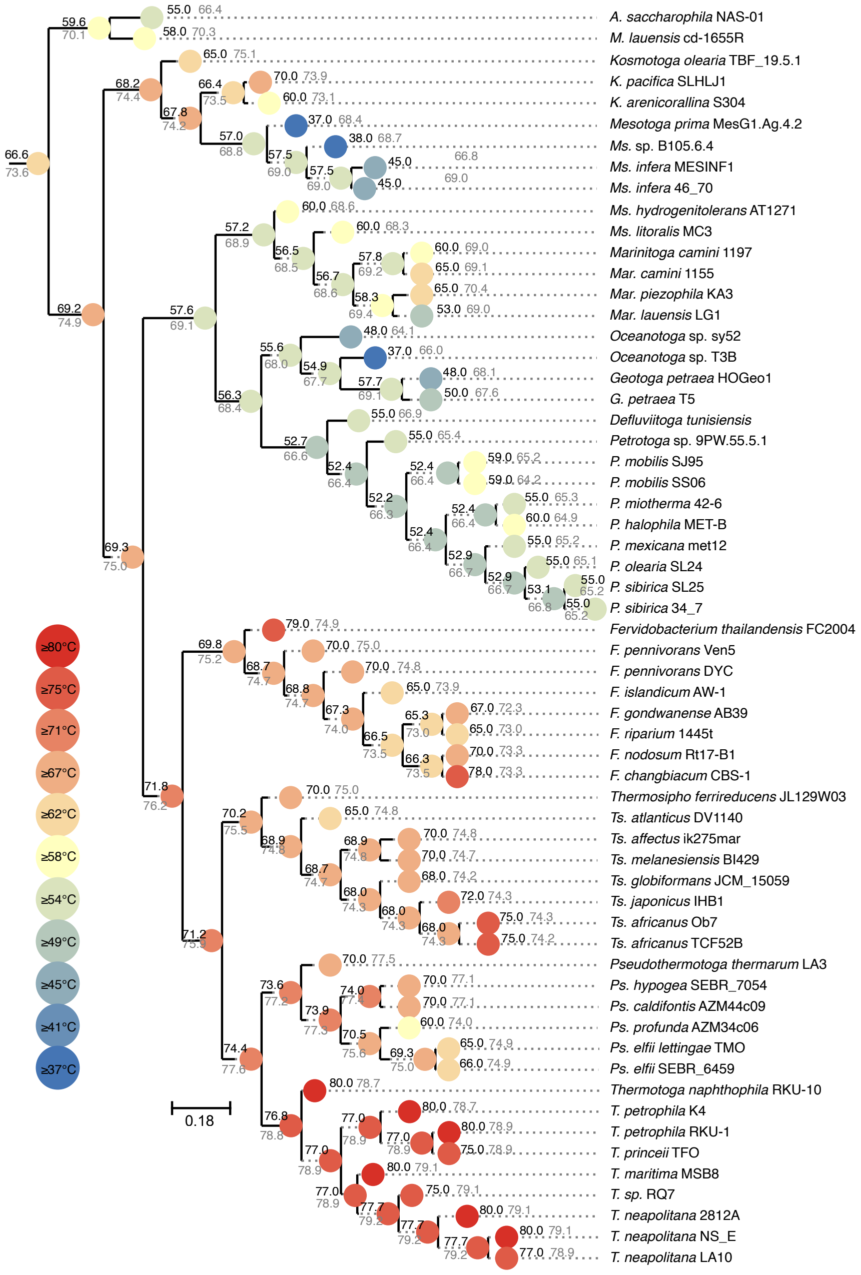

**Supplemental Figure S4. Presence of 29 gene families that correlates with OGT of 37-45°C.** Each row represents a family (HOG#) and each column corresponds to one of the 64 *Thermotogota* species, arranged according to the reference phylogeny (shown on top). Alongside each taxon name is its OGT in degrees Celsius and as a color square. For each gene family, presence of the gene in a taxon is indicated by a square colored by the OGT of the taxon. Additional information of each gene family, including its functional annotation, is provided in **Supplementary Table S3**.

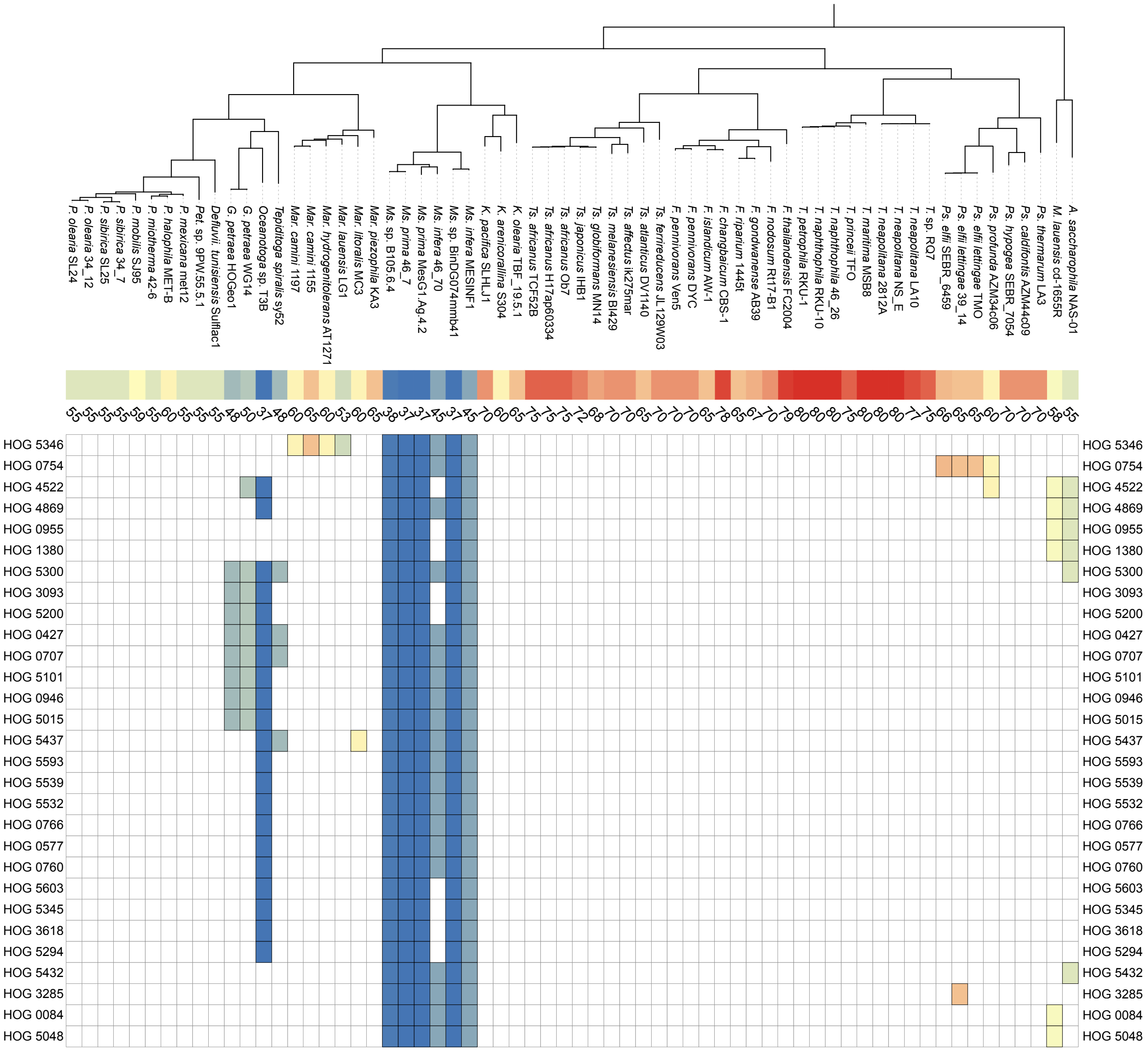

**Supplemental Figure S5. Presence of 9 gene families that correlates with OGT of 48-59°C.** Each row represents a family (HOG#) and each column corresponds to one of the 64 *Thermotogota* species, arranged according to the reference phylogeny (shown on top). Alongside each taxon name is its OGT in degrees Celsius and as a color square. For each gene family, presence of the gene in a taxon is indicated by a square colored by the OGT of the taxon. Additional information of each gene family, including its functional annotation, is provided in **Supplementary Table S4**.

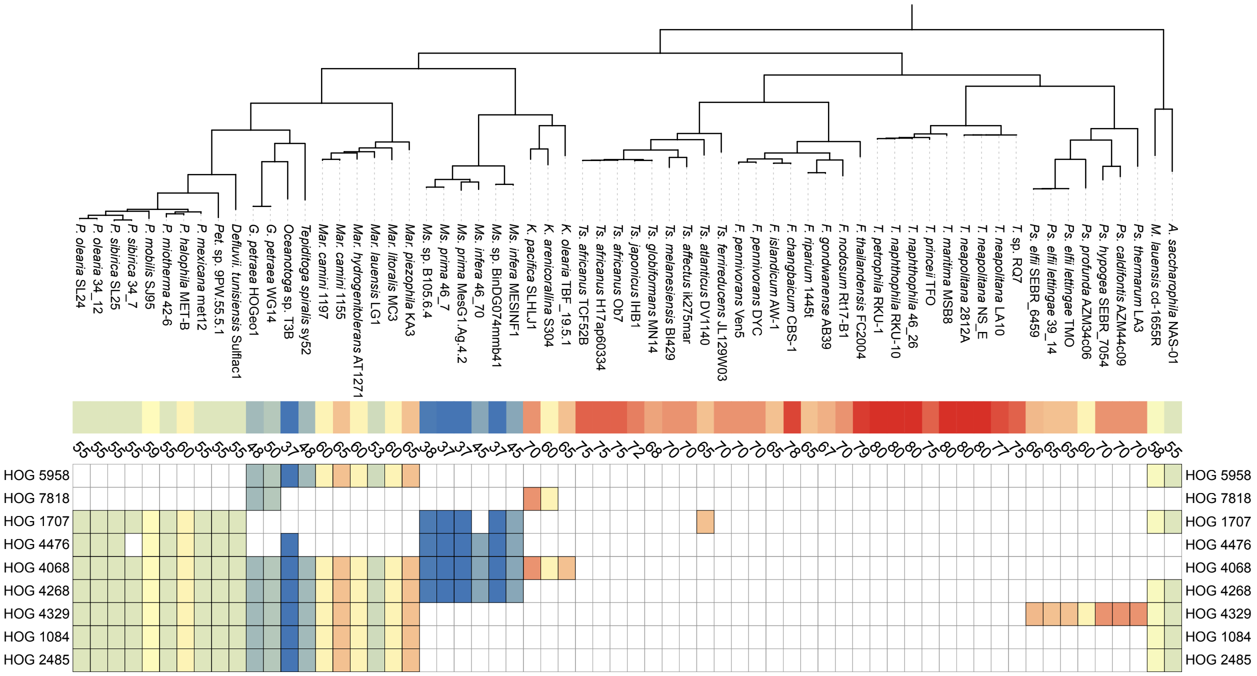

**Supplemental Figure S6. Presence of 30 gene families that correlates with OGT of 60-80°C.** Each row represents a family (HOG#) and each column corresponds to one of the 64 *Thermotogota* species, arranged according to the reference phylogeny (shown on top). Alongside each taxon name is its OGT in degrees Celsius and as a color square. For each gene family, presence of the gene in a taxon is indicated by a square colored by the OGT of the taxon. Additional information of each gene family, including its functional annotation, is provided in **Supplementary Table S5**.

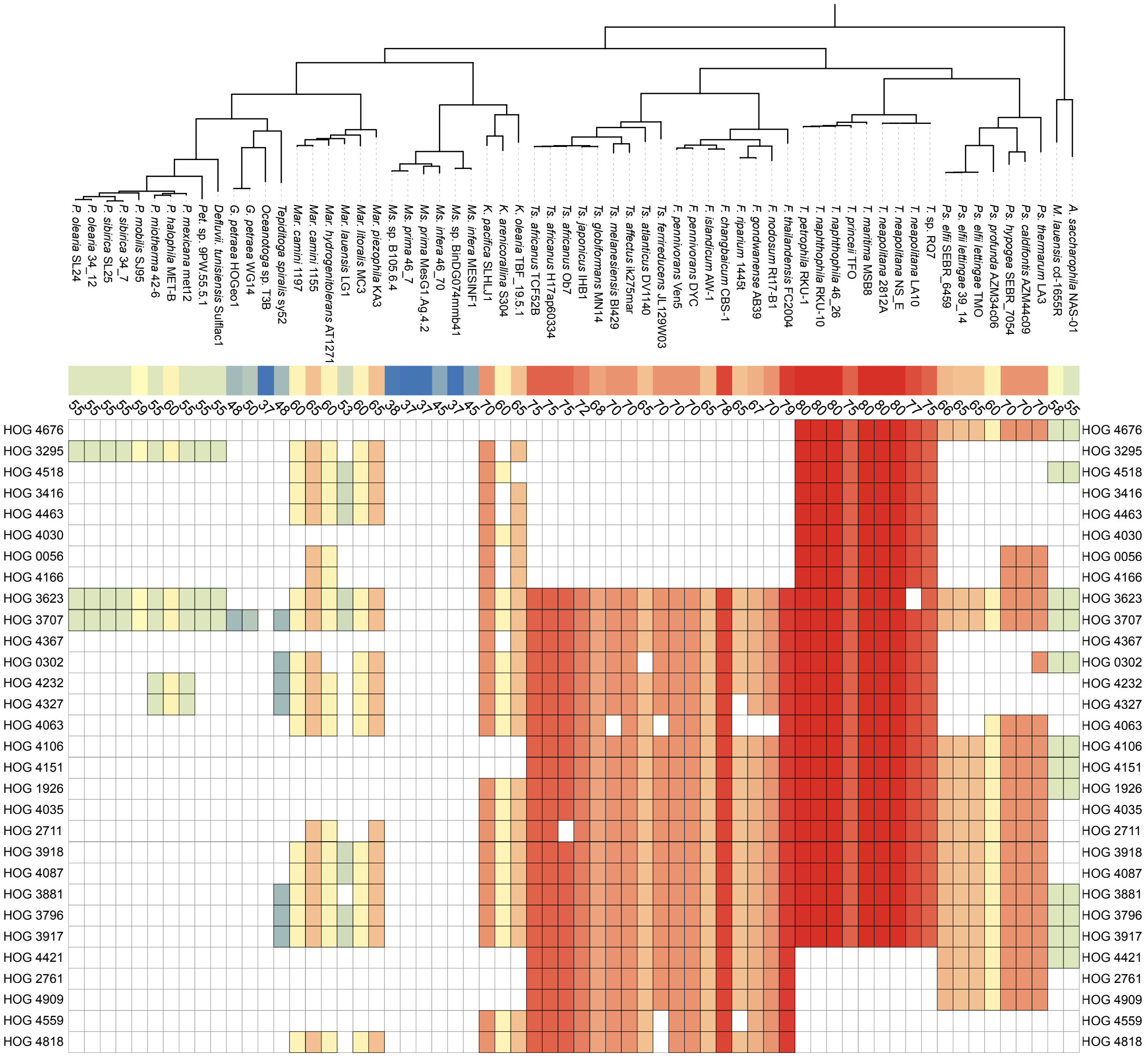

**Supplemental Figure S7. Inferred origins of the 68 gene families in the *Thermotogota* phylum.** The origins are mapped onto a rooted reference phylogeny (outgroup not shown). Each star contains the number of gene families that are inferred to originate along the branch. Stars are colored by the three OGT intervals: blue (37-45°C), green (48-59°C) and red (60-80°C). Stars are scaled with respect to the number of gene families they represent. Scale bar, substitutions per site.

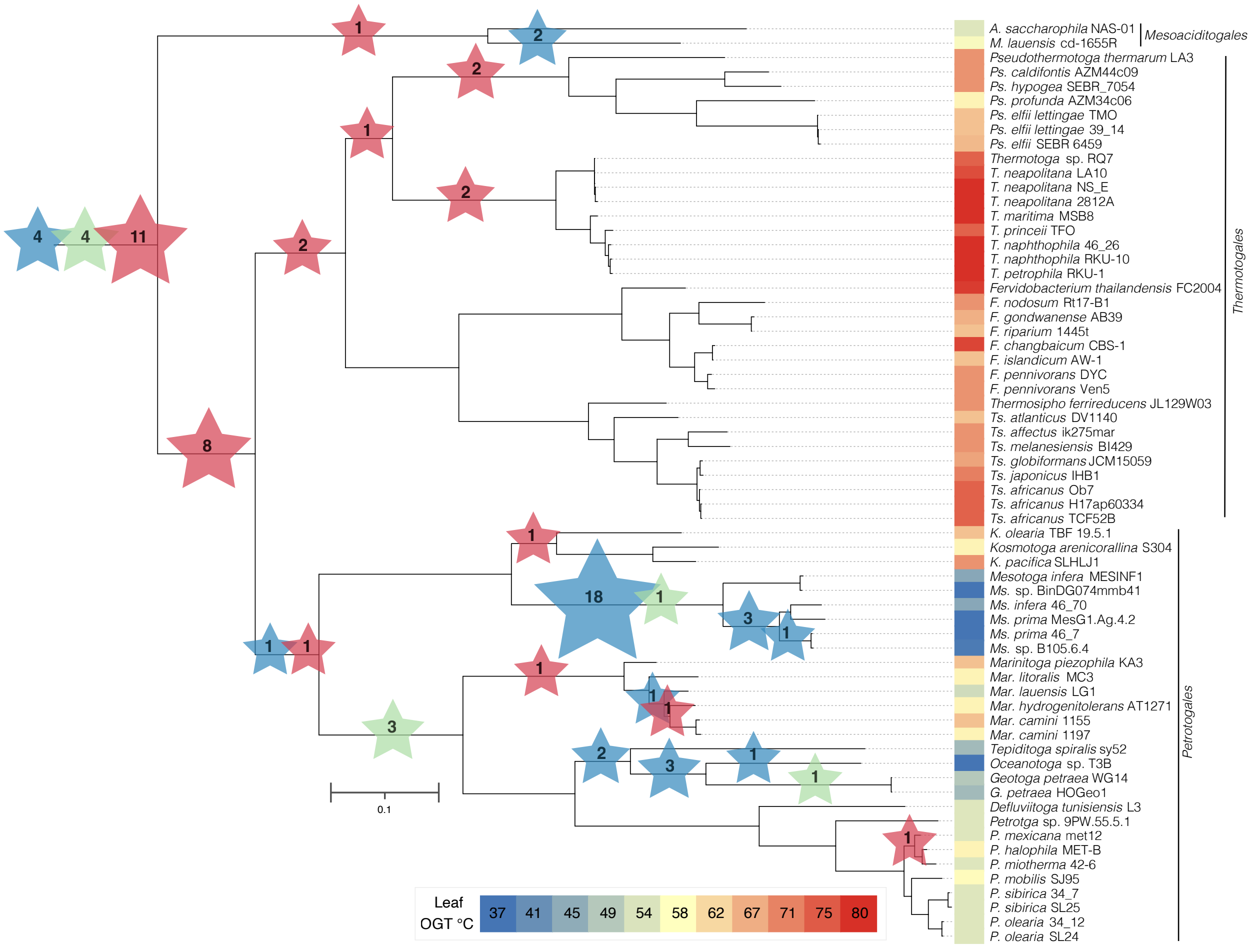

**Supplemental Figure S8. Predicted cellular localization of the 68 OGT-associated gene families. Panel A.** Summary of the total number of gene families (in circles) in each cellular location. **Panel B.** The distribution of gene families' cellular localizations across the three groups with different OGT. Color of each bar corresponds to the localizations shown in **Panel A**.

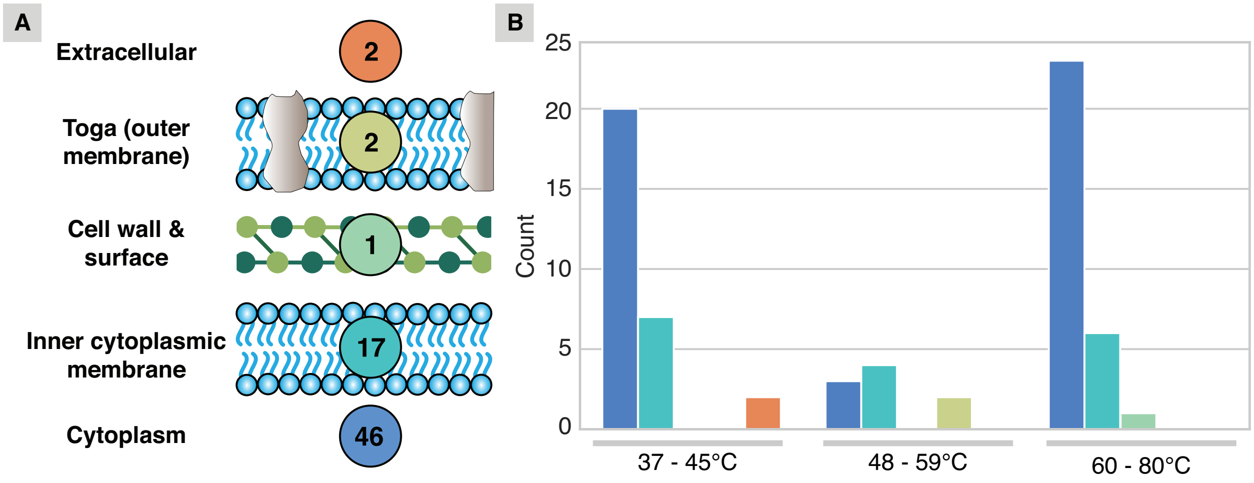

**Supplemental Figure S9. Expanded phylogenies for 19 of the 23 gene families, whose presence correlates best with growth at one of the three temperature intervals.** The phylogenies of 4 other families are shown in **Figure 6**. For each gene family (labeled by the HOG #), its unrooted phylogeny is shown. *Thermotogota* taxa within the gene family are represented by green circles and labeled with genus names. Additional *Thermotogota* homologs, likely from paralogous gene families, are marked by blue circles. Selected sister taxa to *Thermotogota* are also labeled with genus names. Some taxa that are very distantly related to the gene family are collapsed into triangles. "HOG#" labels of gene families associated with OGT of 30-45°C are shown in blue, while the ones associated with OGT of 60-80°C are shown in red. Additional information about these gene families can be found in **Table 1**. Fully labeled phylogenies in Newick and PDF formats are available in the **FigShare** repository.

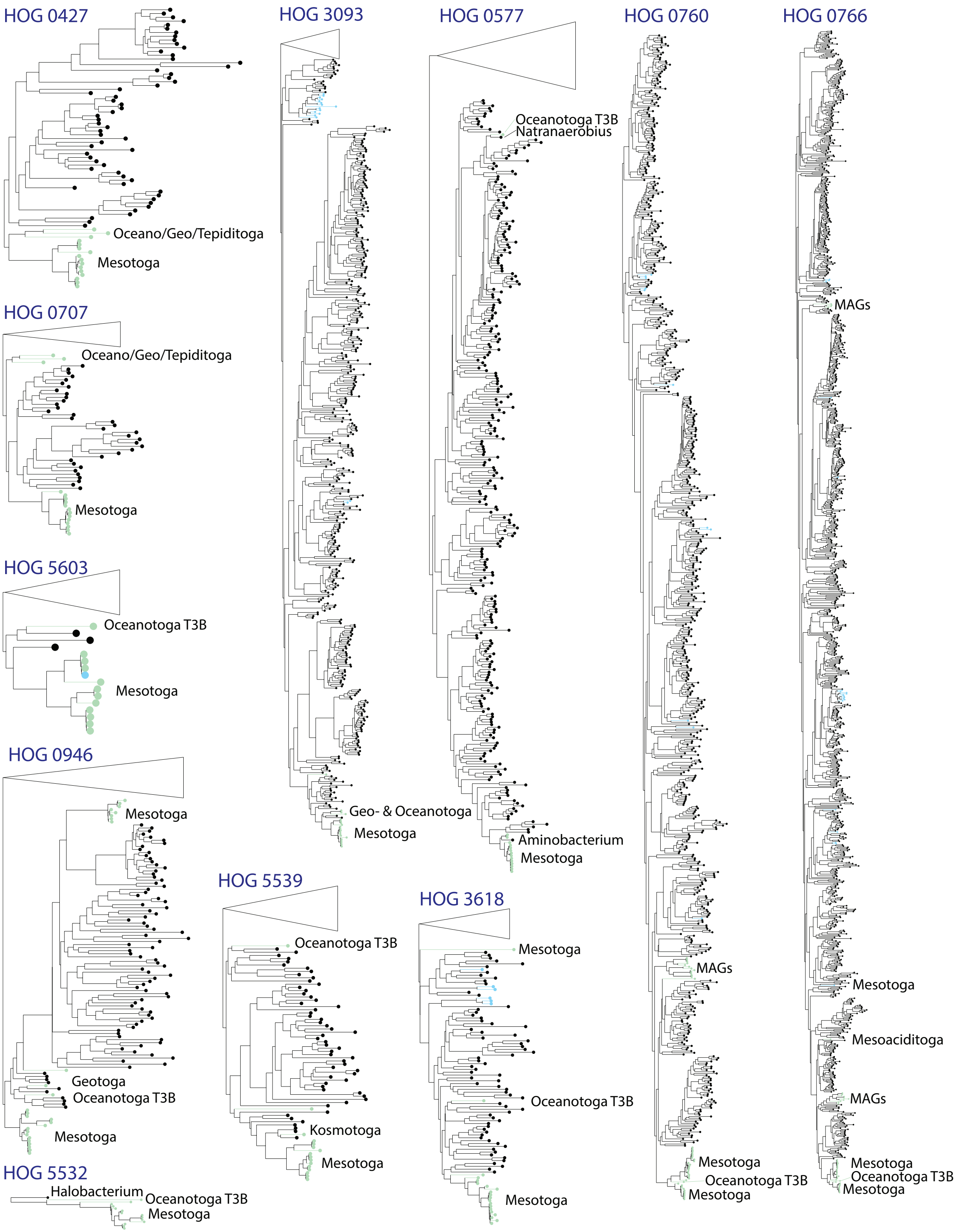
HOG 0427
Oceano/Geo/Tepiditoga
Mesotoga
HOG 3093
Geo- & Oceanotoga
Mesotoga
HOG 0577
Oceanotoga T3B
Natranaerobius
Aminobacterium
Mesotoga
HOG 0760
MAGs
Mesotoga
Oceanotoga T3B
Mesotoga
HOG 0766
MAGs
Mesotoga
Mesoaciditoga
MAGs
Mesotoga
Oceanotoga T3B
Mesotoga
HOG 0707
Oceano/Geo/Tepiditoga
Mesotoga
HOG 5603
Oceanotoga T3B
Mesotoga
HOG 0946
Mesotoga
Geotoga
Oceanotoga T3B
Mesotoga
HOG 5532
Halobacterium
Oceanotoga T3B
Mesotoga
HOG 5539
Oceanotoga T3B
Kosmotoga
Mesotoga
HOG 3618
Mesotoga
Oceanotoga T3B
Mesotoga

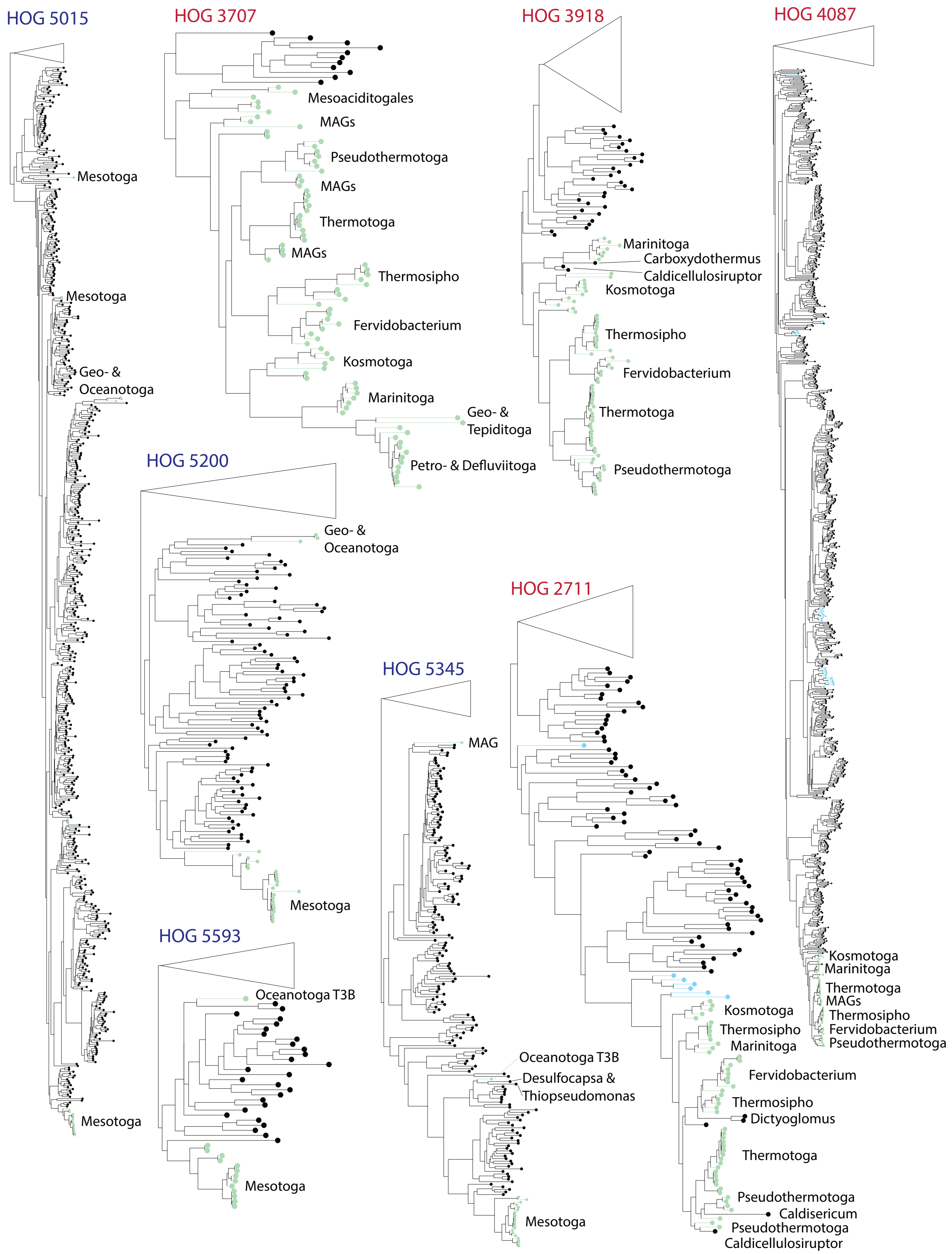

HOG 5015
Mesotoga
Mesotoga
Geo- & Oceanotoga
Mesotoga
HOG 3707
Mesoaciditogales
MAGs
Pseudothermotoga
MAGs
Thermotoga
MAGs
Thermosipho
Fervidobacterium
Kosmotoga
Marinitoga
Geo- & Tepiditoga
Petro- & Defluviitoga
HOG 3918
Marinitoga
Carboxydothermus
Caldicellulosiruptor
Kosmotoga
Thermosipho
Fervidobacterium
Thermotoga
Pseudothermotoga
HOG 4087
Kosmotoga
Marinitoga
Thermotoga
MAGs
Thermosipho
Fervidobacterium
Pseudothermotoga
HOG 5200
Geo- & Oceanotoga
Mesotoga
HOG 5593
Oceanotoga T3B
Mesotoga
HOG 5345
MAG
Oceanotoga T3B
Desulfocapsa & Thiopseudomonas
Mesotoga
HOG 2711
Kosmotoga
Thermosipho
Marinitoga
Fervidobacterium
Thermosipho
Dictyoglomus
Thermotoga
Pseudothermotoga
Caldisericum
Pseudothermotoga
Caldicellulosiruptor

**Supplemental Figure S10.** Scree plot from the Pyseer's multi-dimensional scaling analyses of phylogenetic distances.

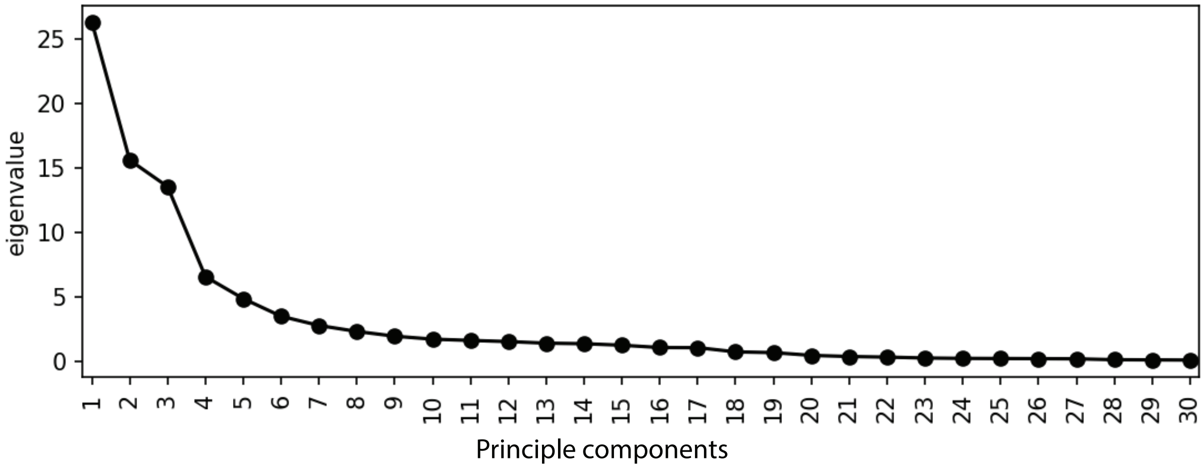
